# Mapping canopy foliar functional traits in a mixed temperate forest using imaging spectroscopy

**DOI:** 10.1101/2024.11.03.621732

**Authors:** Alice Gravel, Etienne Laliberté

## Abstract

- Foliar functional traits are key drivers of ecological processes in forests. Despite progress in forest foliar trait mapping from imaging spectroscopy, there is a need to build environment-specific, spectra-trait models trained from tree-level measurements to improve the accuracy of local trait maps.
- We mapped 12 foliar functional traits in a mixed temperate forest using airborne imaging spectroscopy. Top-of-canopy foliar samples from tree crowns (n = 166), representing a total of 16 species, were collected using a drone platform to measure foliar traits for individual trees, from which tree-level crown spectra were also determined. Partial least squares regression (PLSR) models were used to predict foliar traits from tree-level reflectance spectra (400-2400 nm).
- These models predicted leaf mass per area (LMA), specific leaf area (SLA) and equivalent water thickness (EWT) with high accuracy (R^2^ > 0.8, %RMSE < 15). Models for pigment, nitrogen and cellulose concentrations showed a moderate performance (R^2^ = 0.53–0.68, %RMSE = 17.24–21.31). Poorest performance was observed for lignin, carbon, leaf dry mass content (LDMC) and hemicellulose (R^2^ = 0.24–0.44, %RMSE = 20.67–26.13).
- High-resolution (1.25 m pixel^-1^) foliar trait maps were produced for the entire 16-km^2^ study area. Our study adds to the extensive research aiming to use remote sensing to monitor forest functional trait biodiversity at larger scales and provides models that capture intraspecific variation across many tree species from a mixed temperate forest in eastern Canada.

## Introduction

Biological diversity on Earth has been altered by human activities at an increasing rate over the past century (Chapin III et al., 2000). Changes in plant biodiversity can impact ecosystem processes and their resilience to perturbations (Chapin III et al., 2000; Linke et al., 2006). To monitor these changes, it is important to improve methods for remote sensing of plant biodiversity at large spatial scales. Monitoring plant functional biodiversity, i.e. the diversity in plant functional traits, has attracted particular attention (Cavender-Bares et al., 2020; Jetz et al., 2016) because traits help explain species responses to environmental changes, and traits also control ecosystem functioning (Wang et al., 2020). Among plant functional traits, canopy foliar traits are important because they are key drivers of ecological processes such as primary production, litter decomposition, and nutrient cycling (Curran, 1989). Remote sensing of plant functional traits can enhance our understanding of ecosystem functioning, as well as vegetation responses to climate change (Martin et al., 2018).

Functional traits are morphological, physiological or phenological characteristics that influence plant fitness. They determine growth, reproduction and survival which are then translated into an organism’s performance (Violle et al., 2007). At the leaf level, this includes foliar properties related to photosynthesis (chlorophylls, nitrogen), defense (phenols) and structure (cellulose, lignin; Jacquemoud & Ustin, 2019; Jetz et al., 2016). For example, leaf mass per area (LMA), the ratio of a leaf’s oven-dry mass to its one-sided area, and its inverse, specific leaf area (SLA), represent a trade-off in resource acquisition, reflecting different allocation strategies in construction and maintenance of leaves. This leaf economic spectrum (LES) ranges from fast-productive species (called “acquisitive”), to slow-persistent species (“conservative”) (Cornelissen et al., 2003; Wright et al., 2004). In addition, foliar concentrations of nitrogen and chlorophylls correlate with photosynthetic productivity, and are linked to primary production (Blackburn, 2006). As for structural traits, lignin plays a role in the decomposition of leaves and quality of organic soil. Like cellulose, the amount of lignin provides clues to the quality of the environment for plant growth, but also to nutrient cycling rates in the soil (Jacquemoud & Ustin, 2019; Kokaly et al., 2009). Mineral nutrients are also foliar traits of interest, as they define general metabolic processes by carrying out various specific functions. For example, magnesium is a constituent of chlorophyll, making it an essential element to photosynthesis. Knowledge of their quantity therefore informs the growth and nutritional state of plants (Jacquemoud & Ustin, 2019; Martin, 2020).

Initiatives have been developed to build traits databases, such as TRY (Kattge et al., 2011). However, available trait data covers about only 2% of known vascular plant species on the planet (Jetz et al., 2016). There is a need to fill these trait data gaps, but also to enhance the spatial scale at which functional trait data are acquired. Field sampling is time-consuming and often limited to small areas, and complementary remote sensing approaches are required to traditional field sampling (Linke et al., 2006; Wang et al., 2020).

Imaging spectroscopy, or hyperspectral imagery, has been the leading method providing estimation of foliar traits at the landscape level (Martin et al., 2018). Imaging spectrometers measure reflected solar radiation in hundreds of narrow spectral bands, leading to a complete visible-to-shortwave infrared reflectance spectrum (VSWIR, 400–2500 nm; Asner & Martin, 2009; Goetz et al., 1985). Reflectance of foliage is controlled by its chemical or physical components, including many functional traits of interest in plant ecology (Croft & Chen, 2018). Curran (1989) showed that multiple absorption features in the VSWIR are related to leaf chemicals. Much research has focused on mapping functional foliar traits especially in tropical and mixed temperate forests, and generally on a small number of traits (Serbin et al., 2014; Singh et al., 2015; Zhang et al., 2008). Recently, more studies have successfully retrieved many different foliar traits at the landscape level using imaging spectroscopy (Martin et al., 2018; Wang et al., 2020).

The task of trait mapping requires a direct link between functional traits and spectral signatures, both in time and space (Serbin & Townsend, 2020). To do so, different approaches have been explored, especially partial least square regression (PLSR) modeling (Gara et al., 2019). With this machine-learning method, foliar samples from tree crowns are first collected to measure functional traits. Then, the in-situ measurements are linked to the spectra of pixels corresponding to the sample locations (Serbin & Townsend, 2020). Plot-level estimates of traits can be calculated, e.g., as a weighted average per trait (Wang et al., 2020). This method minimizes the impact of potential errors in geopositioning between the field-collected samples used for trait analyses and imaging spectroscopy pixels, but it also averages out any variation that may occur among and within species in the plot. A different approach was employed in this study. Before field sampling, all trees were first mapped, segmented and identified at the species level from high-resolution drone imagery, building from the work of Cloutier et al. (2023). The top-center part of each tree was sampled with an RTK drone leaf sampling system (Charron et al., 2020; Schweiger et al., 2020), allowing us to precisely target the crown from each selected tree. Then, imaging spectroscopy was aligned with the georeferenced trees to precisely link a pixel to each sampled tree. This approach not only accounts for spectral differences between the crown periphery and center (Schweiger et al., 2020) but also helps the development of models that better consider inter- and intra-specific trait variation, which can improve predictions of PLSR models (Burnett et al., 2021).

In this study, we aimed to map 12 foliar traits at the canopy-level with an approach combining imaging spectroscopy and machine learning. To achieve this, we linked hyperspectral imagery data and field measurements to develop PLSR models for foliar traits retrieval. We mapped predictions of foliar functional traits across the entire study area, as well as their uncertainties, and evaluated the confidence in our trait maps. Based on results from previous studies (Meerdink et al., 2016; Singh et al., 2015; Wang et al., 2019), we hypothesized that : (1) leaf mass per area (LMA), equivalent water thickness (EWT) and cellulose would be predicted with the highest accuracy and (2) most important wavelengths for estimating traits would be in agreement with known spectral absorption characteristics for these traits.

## Material and methods

### Study site

From July 11 to 24, 2022, a field campaign was conducted in a mixed temperate forest within Université de Montréal’s biological field station (Station de biologie des Laurentides; https://sbl.umontreal.ca/). Covering over 16.4 km^2^, this protected area is located in Saint-Hippolyte (Québec, Canada) between 45°58’ and 46°01’ north latitude, and 73°57’ and 74°01’ west longitude (Savage, 2001). The station is found within the Canadian Shield, where podzolic soils formed over glacial till are predominant. This soil order, distributed under acidic coniferous and mixed forest, is characterized by an accumulation of iron (Fe) and aluminum (Al) in the dark illuvial B horizon (Sanborn et al., 2011). The average annual temperature is 3 °C with an annual rainfall between 900 to 1100 mm. The terrain features mounds, rocky escarpments and hills, with an altitude varying from 270 to 450 m above sea level. The site holds many small lakes, streams and wetlands. Disturbances like fires and logging have occurred in the last century, with the most recent forest fire estimated to have occurred around 1923 (Savage, 2001). Since the creation of the research station in 1963, there has been considerable biological and ecological research in the area. The site is covered by a diverse range of broadleaf and evergreen conifer tree species, including *Abies balsamea* (Linnaeus) Miller, *Acer rubrum* Linnaeus, *Acer saccharum* Marshall, *Betula papyrifera* Marshall, *Populus grandidentata* Michaux and *Thuja occidentalis* Linnaeus (Savage, 2001).

### Field sampling

We collected fully sunlit foliar samples from tree crowns using an unoccupied aerial system (UAS, <u>Matrice 300 RTK</u>, Shenzhen, China) equipped with the <u>DeLeaves</u> sampling tool (Charron et al., 2020; Schweiger et al., 2020; Sherbrooke, Quebec, Canada). We sampled 170 trees in a ∼50 ha area surrounding Lac Croche (Fig. 1). We sampled a total of 16 tree species, which together represents most of the species commonly found in the area. Trees in this study included balsam fir (*Abies balsamea*), maples (*Acer pensylvanicum* Linnaeus, *Acer rubrum*, *Acer saccharum*), birches (*Betula alleghaniensis* Britton, *Betula papyrifera*), beech (*Fagus grandifolia* Ehrhart), tamarack (*Larix laricina* [Du Roi] K. Koch), white pine (*Pinus strobus* Linnaeus), poplars (*Populus grandidentata*, *Populus tremuloides* Michaux), spruces (*Picea* A. Dietrich), white cedar (*Thuja occidentalis*), hemlock (*Tsuga canadensis* [Linnaeus] Carrière) and a few less common species (red oak; *Quercus rubra* Linnaeus, ironwood; *Ostrya virginiana* [Miller] K. Koch). Black and red spruces often hybridize naturally when they grow together (Perron & Bousquet, 1997), making the identification to the species level difficult. For this reason, all spruces were grouped together by genus. Aside from tamarack crowns that were clustered in a bog, each species was distributed somewhat evenly over the sampling area (Fig. 1). We used accurately georeferenced delineated tree crowns from Cloutier et al. (2023) to select tall trees with crowns that had a diameter greater than 1 meter for sampling. This allowed us to optimize the drone platform use (visibility for the operator) as well as guarantee the feasibility of linking trait data to spectral data.

**Figure 1.**
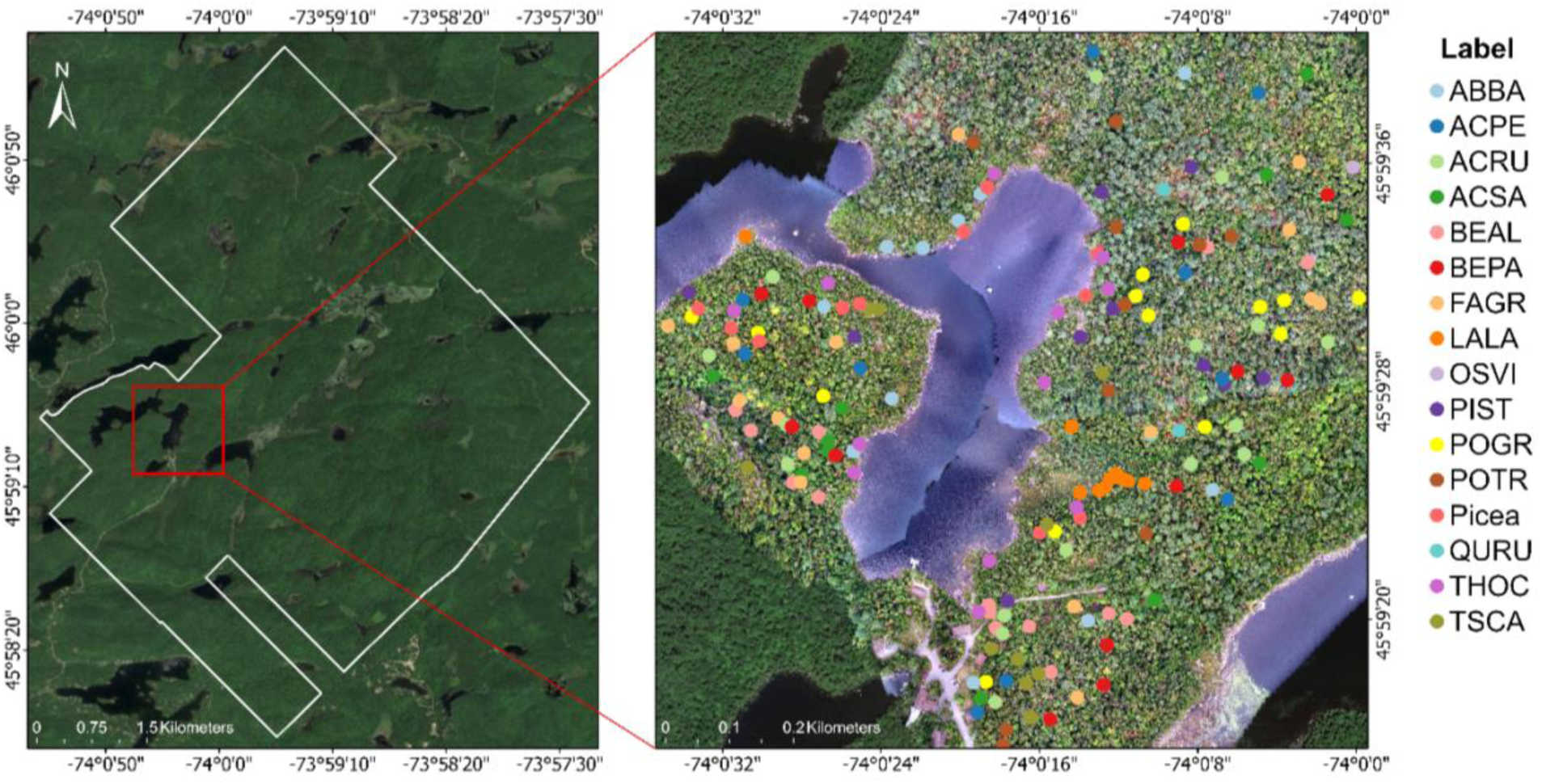
Map of the sampling area within the study site, Saint-Hippolyte, Québec, Canada. The left panel shows the station’s boundaries in white. The right panel zooms on the sampling area, which shows the sampled trees, with each color referring to a different tree species. Only spruces were grouped by genus. ABBA, *Abies balsamea*; ACPE, *Acer pensylvanicum*; ACRU, *Acer rubrum*; ACSA, *Acer saccharum*; BEAL, *Betula alleghaniensis*; BEPA, *Betula papyrifera*; FAGR, *Fagus grandifolia*; LALA, *Larix laricina*; OSVI, *Ostrya virginiana*; PIST, *Pinus strobus*; POGR, *Populus grandidentata*; POTR, *Populus tremuloides*; Picea, *Picea* sp.; QURU, *Quercus rubra*; THOC, *Thuja occidentalis*; TSCA, *Tsuga canadensis*.

### Trait analyses

The UAS-collected leaf samples were stored in sealed plastic bags a few minutes after collection. A first subset of the samples was weighed for fresh mass, rehydrated for 6 hours, weighed again, and scanned for total rehydrated leaf area. Then, leaves were oven-dried at 65°C for 72h and weighed for total dry mass. This workflow followed the protocol from Laliberté (2018) to determine specific leaf area (SLA, m^2^ kg^-1^), leaf mass per area (LMA, g m^-^ _2_), leaf dry matter content (LDMC, mg g^-1^) and equivalent water thickness (EWT, mm). A second subset of the samples were used to obtain fresh leaf disks that were frozen to -80 °C. Chlorophylls (*a* and *b*, in mg g^-1^) and bulk carotenoids (mg g^-1^) were measured by spectrophotometry using a plate reader (Girard et al., 2021; <u>SPECTROstar Nano</u>, Ortenberg, Germany). The remaining subset of leaves was immediately oven-dried at 65 °C. These dried leaves were then ground for further chemistry analysis. Leaf carbon fractions (hemicellulose, cellulose, and lignin; %) were determined using a sequential digestion (<u>ANKOM2000 Fiber</u> <u>Analyzer</u>, Macedon, NY, USA) following Ayotte & Laliberté (2019). Carbon and nitrogen concentrations (C, N; %) were measured with an elemental analyzer (<u>Elementar Vario MICRO</u> <u>Cube</u>, Lyon, France) following Ayotte et al. (2018).

### Imaging spectroscopy data

On July 9^th^, 2022, airborne imaging spectroscopy data was acquired over the study area by a flight crew of the Flight Research Laboratory of the National Research Council Canada (NRC). Imaging spectrometers included both CASI-1500 (CASI, Compact Airborne Spectrographic Imager) and SASI-644 (SASI, Shortwave Airborne Spectrographic Imager). The CASI sensor captures over 288 bands between 375 and 1054 nm, whereas the SASI sensor captures over 160 bands between 883 and 2523 nm. Their respective field of view is 39.9° and 39.7° (Wallis et al., 2023). The spatial resolution of the products was resampled to 1.00 m for CASI and 1.25 m for SASI. Average altitude was 1413 m above sea level and average speed 41.67 m s^-1^. Image preprocessing was done by the Applied Remote Sensing Lab (McGill University, Department of Geography, Montreal, Canada; ARSL) and the Flight Research Laboratory of NRC. The imaging spectroscopy data underwent radiometric, geometric and atmospheric correction. Water absorption bands in the short-wave infrared (SWIR, ∼1550–2500 nm) were removed and interpolated with a straight line during atmospheric correction. This resulted in seven flightlines for each sensor that were analyzed separately. We georeferenced each SASI flightline using a first order polynomial transformation and then aligned CASI data to SASI using an auto-georeferencing tool in ArcGIS Pro version 3.0.1 (ESRI, 2022). The spatial resolution of CASI was down-sampled to that of the SASI data (1.25 m pixels) using a nearest neighbor technique to avoid creating new values and thus minimizing changes in pixel values. All subsequent processing and analyses were conducted in R version 4.2.1 (R Core Team, 2022).

From the georeferenced delineated tree crowns (Cloutier et al., 2023), we created a 0.5 m inner buffer for each field-sampled tree crown. We extracted CASI and SASI pixels within those buffers to obtain mean spectral reflectance per tree crown (Fig. S1). The buffer minimized the chance that neighboring pixels from other tree crowns contributed significantly to the calculation of the average reflectance. This was also done because the ground instantaneous field of view of a sensor is often larger than the pixel size, resulting in potential pixel overlap (Jones & Vaughan, 2010). Thus, the buffer allowed to minimize this contamination effect possible from the adjoining pixels. We applied an NDVI filter (>0.6) and a shadow threshold at 802 nm (>0.10) to remove non vegetated and/or shaded pixels. We removed low signal and noisy regions in CASI (375–395 nm, 1000–1060 nm) and SASI (2400–2442 nm). Both sensors were matched using the R package “spectrolab” (Meireles et al., 2017). To minimize the noisy signal without hiding absorption features, we applied a conservative smoothing on mean spectra using the R package “hsdar” (Lehnert et al., 2019) with a window size of 3 and a 2^nd^ polynomial order (Fig. S2). Smoothed spectra were vector-normalized to reduce inner canopy shade and brightness differences between flightlines (Fig. S3). Four tree crowns with insufficient signal in pixels were removed, resulting in 166 crowns for model development. Summary statistics of foliar traits for these crowns are available in Table S1.

### Linking foliar traits to spectral data

We used partial least square regression (PLSR) to estimate leaf traits from VSWIR reflectance. This machine learning technique is widely used for estimating foliar properties from spectral reflectance data (Singh et al., 2015; Wang et al., 2020). During the calibration phase, the predictors, here the spectral reflectance values at different wavelengths, are linked to the observations, which are the foliar traits (Schweiger et al., 2020). PLSR modeling handles the high collinearity between predictors and the larger number of predictors compared to observations. The wavelengths are reduced to a small number of uncorrelated components or latent variables that are maximally correlated with the traits (Burnett et al., 2021).

For each foliar trait, an ensemble of PLSR models was developed, following Burnett et al. (2021) and Kothari et al. (2023). The entire spectral range (400–2400 nm) was used and no transformation was applied to the trait dataset. Interpolated water absorption bands were included in the analysis since it did not show major differences in model development to remove them. We first split the data into calibration (30%) and validation (70%) subsets. Initial models were built using a 10-fold cross-validation on the calibration subset, to determine the optimal number of components while avoiding overfitting. We selected the smallest number of components that showed a root mean squared error of prediction (RMSEP) within one standard error of the global minimum. However, in the case of LDMC, hemicellulose, cellulose, chlorophyll *b* and carbon, a very small number of components was suggested using this approach (e.g., 1 or 2). Therefore, for these traits, we selected a number of components that showed a low RMSEP (either the absolute minimum or close to the second minimum). This approach allowed us to avoid underfitting of the spectral data by selecting an overly simplistic model that would perform poorly (Burnett et al., 2021). We also calculated the variable influence in projection (VIP) to determine the most significant wavelengths for prediction. The VIP was significant when greater than 0.8 (Singh et al., 2015).

Next, to determine model uncertainties, we performed a jackknife analysis on the calibration subset. We further divided the calibration subset into training (70%) and testing (30%) to conduct a permutation analysis with 100 iterations, resulting in 100 models per trait. The resulting model for each iteration was applied to the 30% testing subset. As internal validation, all models resulting from this resampling procedure were applied to the validation subset, resulting in a distribution of estimates for each sample. The mean estimates were compared to the measured values to determine validation statistics. We evaluated the models’ performance with the coefficient of determination (R^2^), the root mean square error (RMSE), the percent root mean square error (%RMSE) and the bias. To assess general model performance, we focused on the R^2^ that gives insight into the model’s precision, and the %RMSE that indicates the model’s accuracy. The %RMSE is calculated by taking the RMSE between the observed and predicted values, normalized by the range of observed values between the 0.025 and 0.975 quantiles, and expressed as a percentage. With a relative RMSE as a percentage, different models’ accuracy can be compared between them.

### Foliar trait maps

We used the same processing workflow (see section *Imaging spectroscopy data*) for all pixels of the imagery. We applied the ensemble PLSR models (100 iterations) to all pixels, resulting in a mean predicted trait value for each pixel. We mapped the mean predictions and standard deviation (SD) values of the 100 PLSR models to evaluate for uncertainties, following Wang et al. (2020). Because uncertainties are influenced by the range of mean estimates, we also mapped uncertainties in relative terms (coefficient of variation, CV = SD/mean) to compare different trait maps with each other (Verrelst et al., 2016). Pixels whose predictions returned negative values were masked out for analysis (0–1.4% of pixels).

We determined the confidence in the trait predictions by calculating the percentage of pixels with high confidence for each trait and flightline separately, following Wang et al. (2020). First, pixels with unrealistically low or high values were selected with the range of measured traits from the field: trait_min_ - | trait_min_ | × 25% and trait_max_ + | trait_max_ | × 25%. Subsequently, we used a threshold of 25% to identify pixels with unacceptable high relative uncertainties (CV) within the selected range. This filtering resulted in a percentage of pixels with high confidence. We also compared the distribution and the range of measured traits in the field (Table S1) and estimated traits derived from polygons delineating tree crowns covering the sampling area. We used 23 000 annotated tree crowns from Cloutier et al. (2023) and extracted pixels of estimated traits completely within these polygons, resulting in 50 573 pixels. The complete workflow, from field sampling to mapping canopy traits, is summarized in Figure 2.

**Figure 2.**
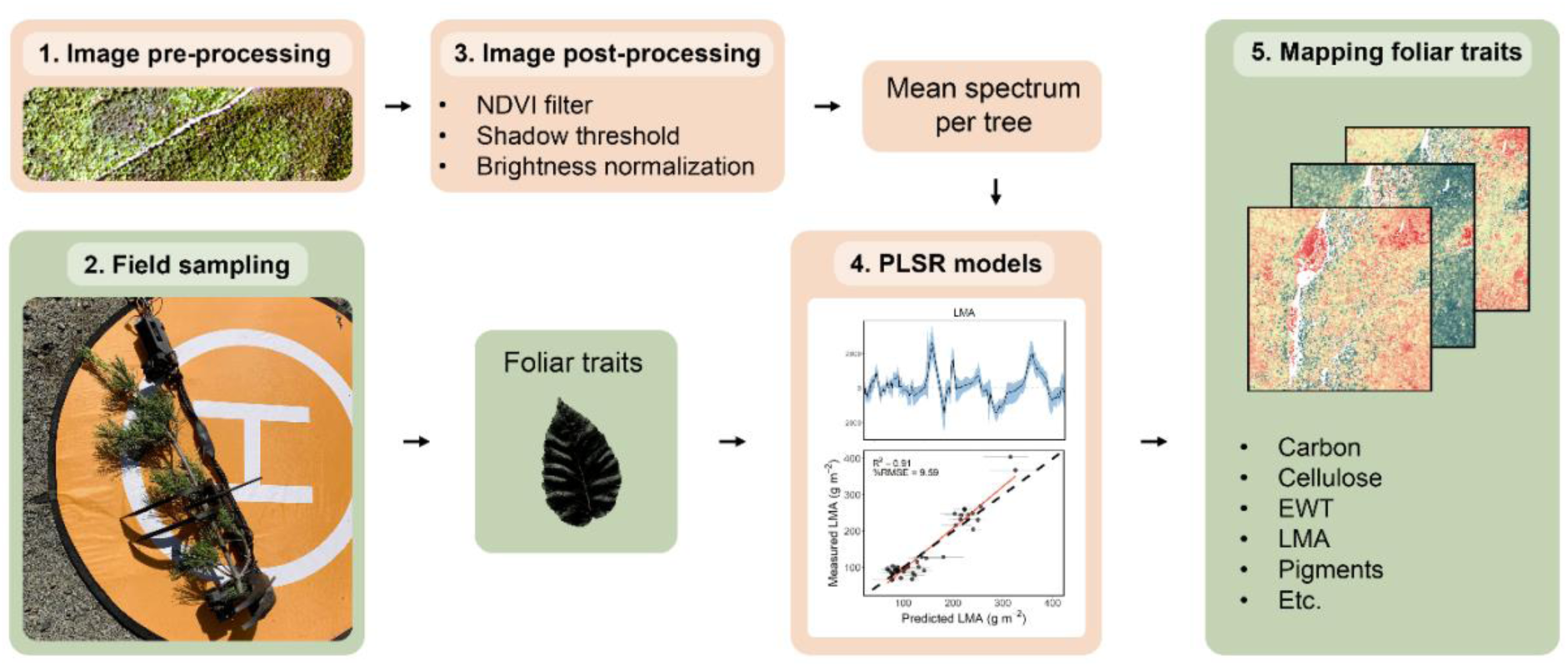
Summarized workflow to map foliar traits. Steps include: 1. Acquire and pre-process imagery with appropriate corrections; 2. Field sampling to acquire trait data; 3. Image post-processing to obtain one mean spectrum per tree crown; 4. PLSR modeling to link spectral data to trait data; 5. Applying ensemble models to imagery to obtain mean estimates per pixel and generate trait maps. This approach assumes a homogeneous trait value within the tree crown.

## Results

### Model performance

The model statistics validation set for each trait is presented in Table 1. For all models, the R^2^ ranged from 0.24 to 0.91, while the %RMSE ranged from 9.59% to 26.13%. Models for LMA, EWT and SLA showed the highest performance, with high R^2^ (>0.8) and %RMSE < 15%. An intermediate performance was obtained for pigments (chl *a*, chl *b* and carotenoids), nitrogen and cellulose, with an R^2^ >0.5 and %RMSE <20 (except for cellulose that showed a %RMSE of 21.31). The poorest model performance was observed for lignin, carbon, LDMC and hemicellulose with an R^2^ <0.5 and a %RMSE >20. Still, all models showed a %RMSE <25, apart from carbon (Table 1). All models tended to overestimate traits, except for LMA and SLA that were under-estimated as shown by the bias (Table 1).

**Table 1.** Validation statistics of foliar functional traits from Partial Least Square Regression (PLSR) models. R^2^, coefficient of determination; RMSE, root mean square error; %RMSE, percent root mean square error. LMA, leaf mass per area; EWT, equivalent water thickness; SLA, specific leaf area; Chl *a*, chlorophyll *a*; Chl *b*, chlorophyll *b*; LDMC, leaf dry matter content. The bias represents the difference between mean predicted and observed values, whereas the components refer to the number of latent vectors used for model building.

| Traits | Units | $R^2$ | RMSE | %RMSE | Bias | Components |
| --- | --- | --- | --- | --- | --- | --- |
| LMA | $\text{g m}^{-2}$ | 0.91 | 28.10 | 9.59 | -0.0797 | 6 |
| EWT | mm | 0.87 | 0.03 | 11.88 | 0.0010 | 7 |
| SLA | $\text{m}^2 \text{kg}^{-1}$ | 0.82 | 1.62 | 13.35 | -0.2086 | 6 |
| Carotenoids | $\text{mg g}^{-1}$ | 0.68 | 0.08 | 17.83 | 0.0033 | 4 |
| Chl *a* | $\text{mg g}^{-1}$ | 0.66 | 0.38 | 17.47 | 0.0879 | 5 |
| Nitrogen | % | 0.65 | 0.30 | 18.42 | 0.0697 | 11 |
| Chl *b* | $\text{mg g}^{-1}$ | 0.64 | 0.12 | 17.24 | 0.0292 | 6 |
| Cellulose | % | 0.53 | 2.04 | 21.31 | 0.4220 | 5 |
| Lignin | % | 0.44 | 2.19 | 20.67 | 0.4448 | 5 |
| Carbon | % | 0.34 | 0.89 | 26.13 | 0.1792 | 7 |
| LDMC | $\text{mg g}^{-1}$ | 0.32 | 39.67 | 23.79 | 2.8146 | 8 |
| Hemicellulose | % | 0.24 | 3.53 | 23.33 | 0.2819 | 5 |

The number of components used for model building ranged from 4 to 11. The nitrogen model had the highest number of components of all traits (11). Figure 3 displays scatter plots comparing predicted traits obtained through internal validation with measured traits from field sampling. These plots provide a visual report of the models fitting. LMA, SLA and EWT models showed the best fit of all traits. Model coefficients are presented in Figure S4. Pigments displayed the largest magnitude of coefficients in the visible spectrum (VIS, ∼400–700 nm), whereas physical, water-related, and structural traits showed the largest magnitude in the SWIR1 (∼1550–1800 nm) or SWIR2 (∼2000–2500 nm).

**Figure 3.**
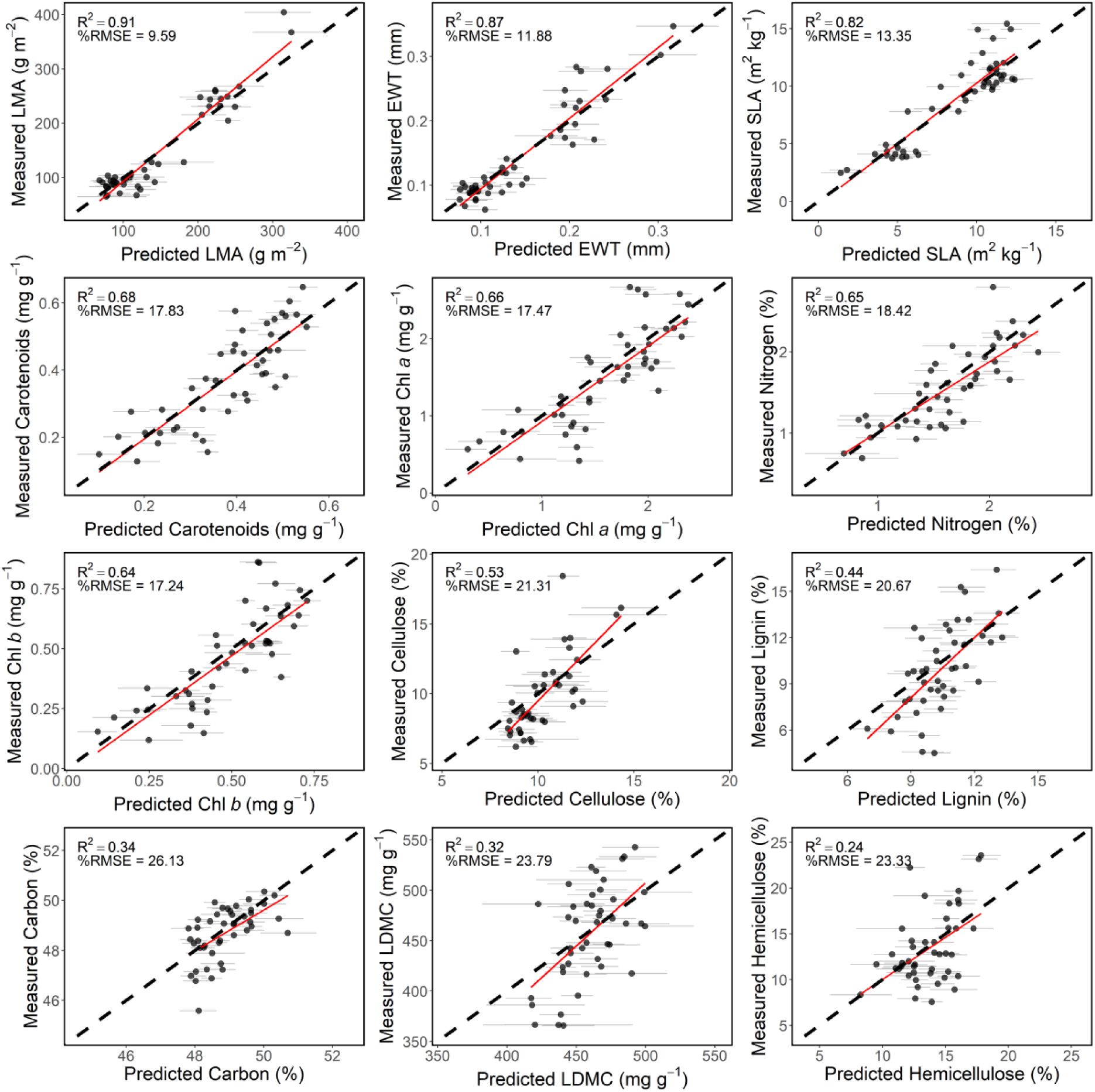
Scatter plots of predicted traits by PLSR models after internal validation vs measured traits from field sampling. Coefficient of determination (R^2^) and percent root mean square error (%RMSE) are shown. Horizontal bars show standard deviation. The dashed black line indicates the 1:1 line and the red line is the fit ordinary least square (OLS) regression between predicted and measured trait values (n = 44). Panels are presented in descending order of R^2^, from top left to bottom right. LMA, leaf mass per area; EWT, equivalent water thickness; SLA, specific leaf area; Chl *a*, chlorophyll *a*; Chl *b*, chlorophyll *b*; LDMC, leaf dry matter content.

### Trait maps and uncertainties

An example of general distribution of four different foliar traits (leaf mass per area, cellulose, chlorophyll *a* and nitrogen) is presented in Figure 4. This area contains a wide variety of conifer (e.g., *Abies balsamea*, *Pinus strobus*, *Thuja occidentalis*) and broadleaf species (e.g., *Acer rubrum*, *Betula papyrifera*, *Populus grandidentata*). Mean trait predictions varied among functional types (conifer vs broadleaf) and species. All mapped traits are presented in Figure S5 and their uncertainties in Figure S6. The mean trait predictions varied along an environmental gradient related to hydrology. Maps of distribution of the same four foliar traits are presented in Figure 5 as an example of trait variation along a gradient of topographic wetness. The area covers a poorly drained forested peatland characterized by higher LMA and cellulose content, while exhibiting lower chlorophyll *a* and nitrogen content (Fig. 5).

**Figure 4.**
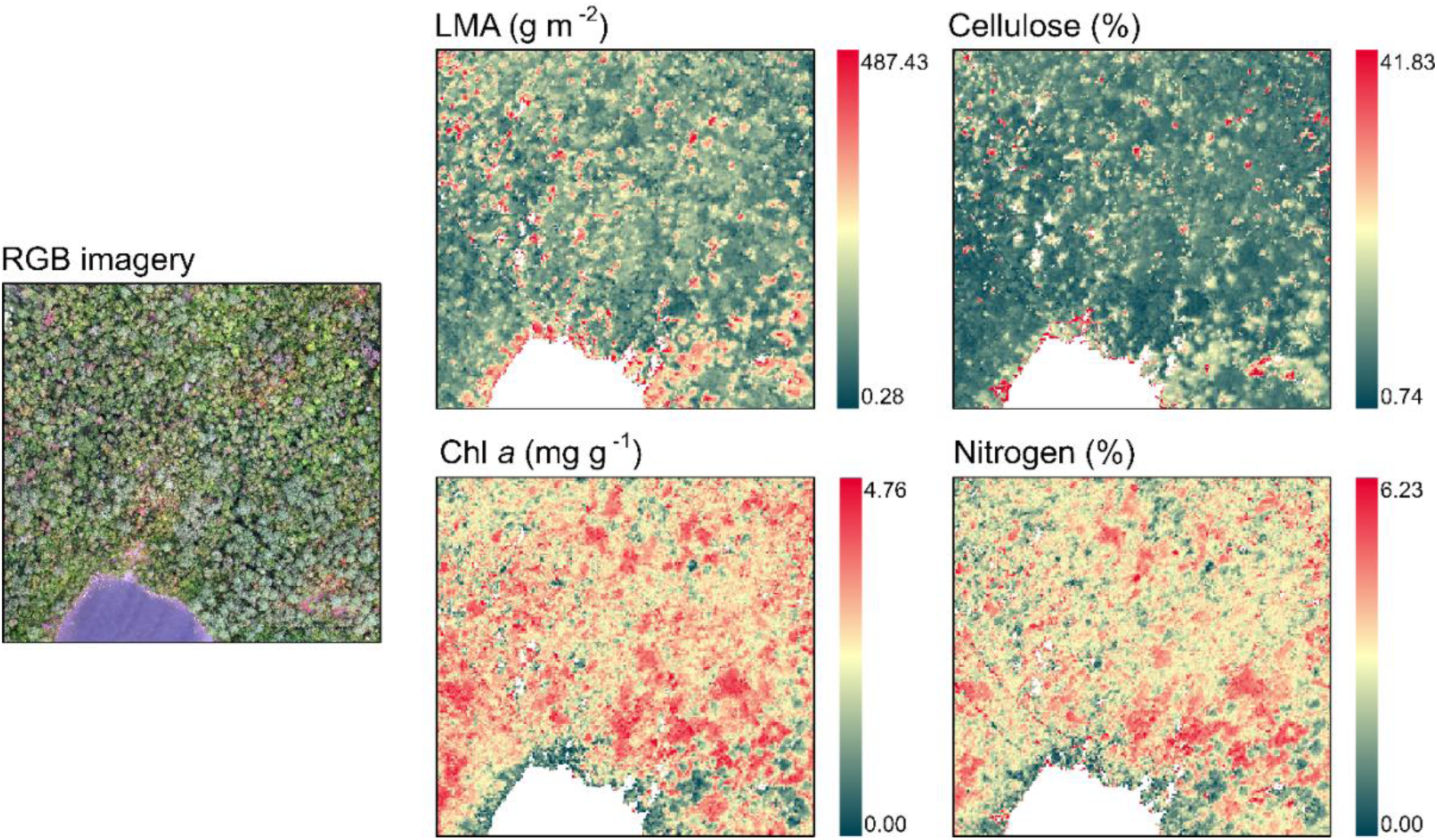
Maps of predicted foliar traits as an example of general distribution of four different foliar traits in the study area (LMA, leaf mass per area; cellulose; Chl *a*, chlorophyll *a* and nitrogen). Trait predictions varies among functional types. RGB imagery was provided by Cloutier et al. (2023).

**Figure 5.**
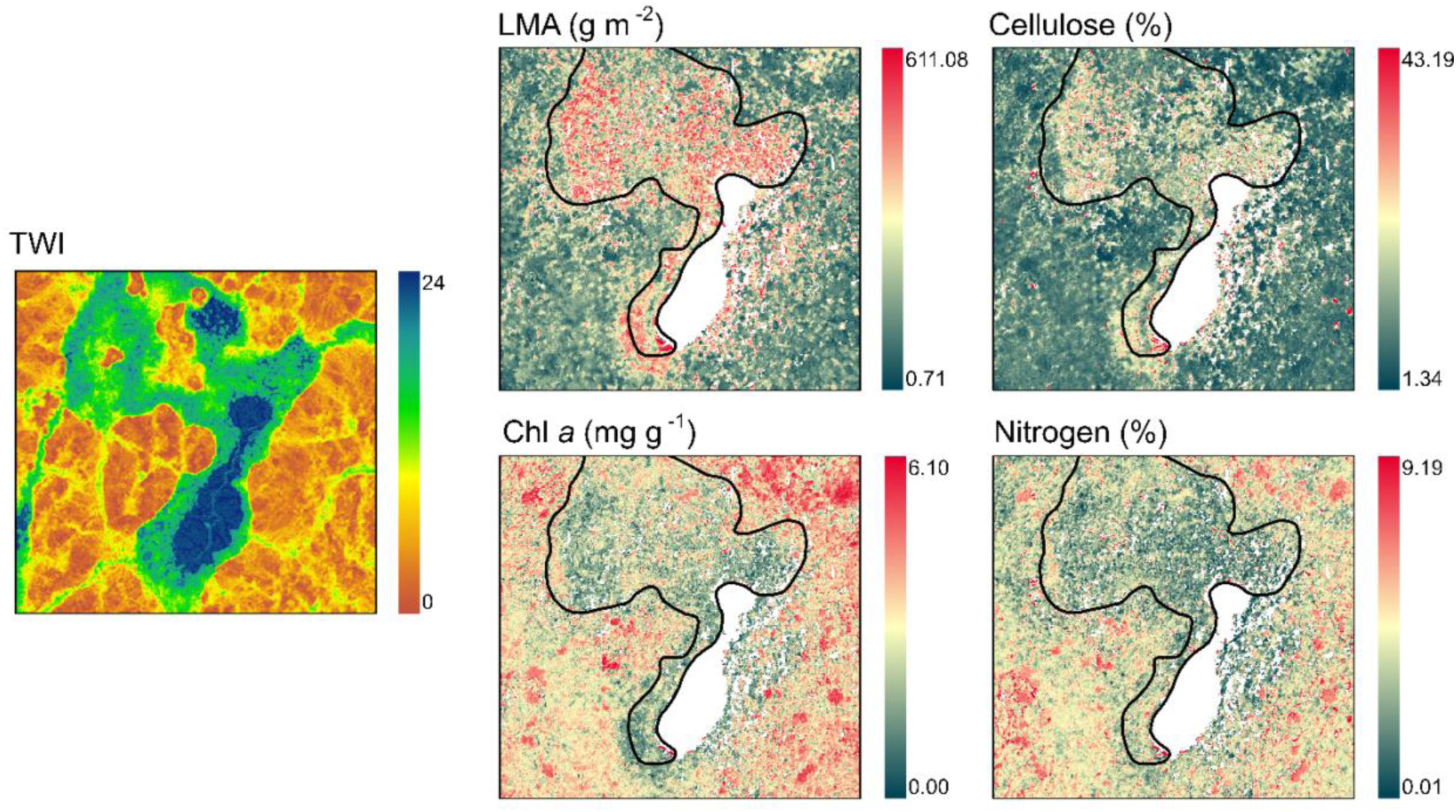
Maps of predicted foliar traits as an example of variation of four different foliar traits along an environmental gradient linked to topographic wetness (LMA, leaf mass per area; cellulose; Chl *a*, chlorophyll *a* and nitrogen). The black polygon represents the location of a forested peatland (Canards Illimités Canada, 2023). TWI : Topographic Wetness Index (Ministère des ressources naturelles et des forêts, 2024). The index increases with the capacity of the land to retain humidity (Ministère des Forêts, de la Faune et des Parcs, 2020).

Trait model uncertainties, which were calculated as the standard deviation of predictions by the set of 100 coefficients, varied among all traits and environmental gradients. The highest uncertainty was observed in wetlands such as swamps, fens and shallow waters, as well as vegetation near lake borders. Figure 6 illustrates an area with shallow waters showing high uncertainties for four different foliar traits. Over 90% of vegetated pixels for all traits were within the predefined range from field measurements (see section *Foliar trait maps*). Among these, over 90% of pixels had relative uncertainties <25% for all traits, except LMA (87% of pixels with high confidence). The percentage of pixels with high confidence for each trait, for all flightlines combined, are shown in Figure S7.

**Figure 6.**
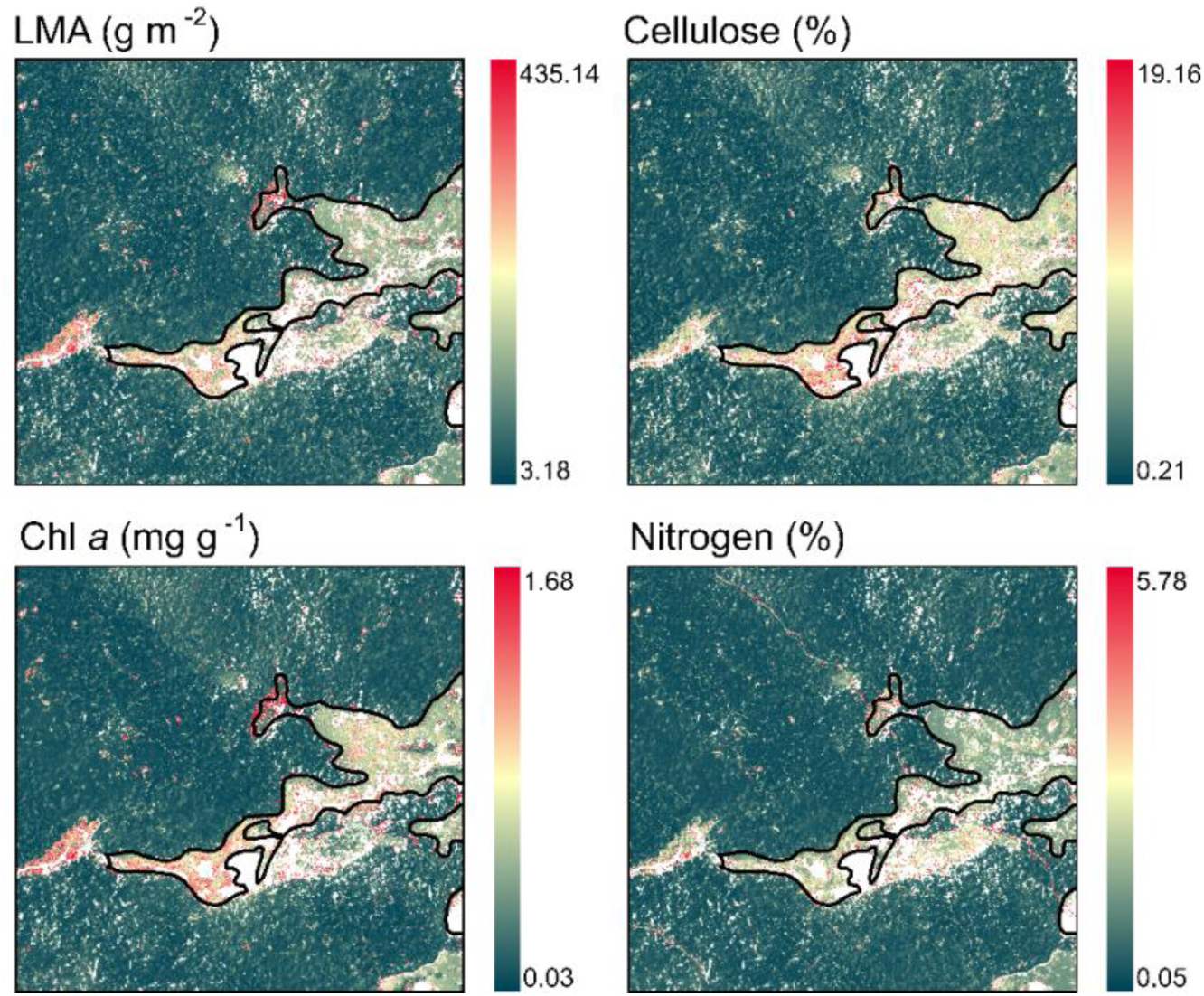
Distribution of model uncertainties for four foliar traits across a wetland (LMA, leaf mass per area; cellulose; Chl *a*, chlorophyll *a* and nitrogen). The black polygon identifies an area of shallow water (Canards Illimités Canada, 2023), corresponding to high uncertainties.

The range of estimated traits mostly covered the range of measured traits from field sampling (Fig. 7). Some traits showed a long right tail of high values (cellulose, EWT, LMA), whereas carbon revealed the opposite with a long left tail of lower values. Overall, there was less variability in the traits predicted by the models, as indicated by the narrow and high peak of measured traits. Also, the standard deviation for estimated traits was lower than measured traits (results not shown).

**Figure 7.**
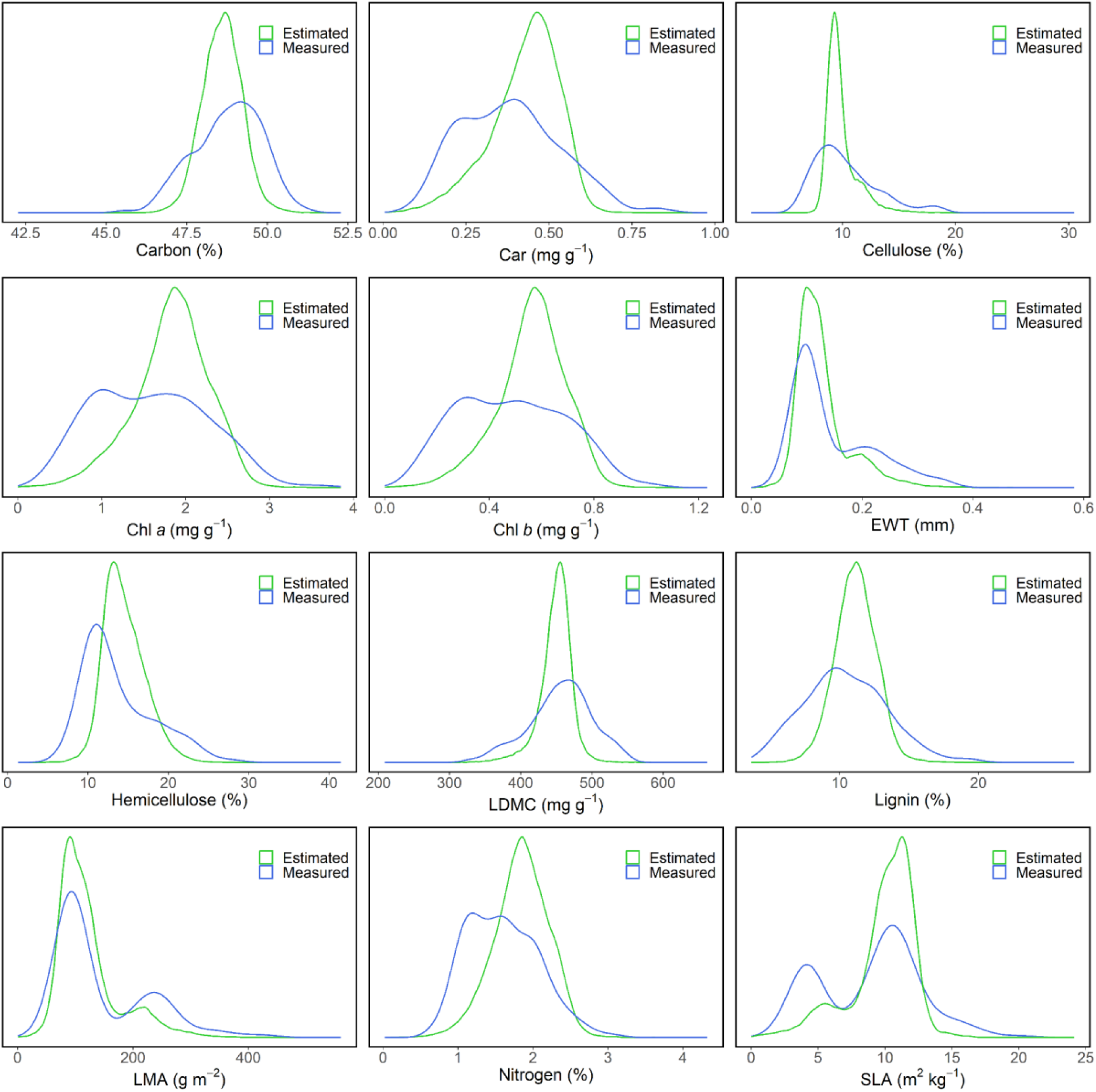
Distribution and range of measured traits in the field (n = 166) and pixels of estimated traits from annotated tree crowns (n = 50 573) (Cloutier et al., 2023). We filtered the dataset to keep labels of interest, removing trees identified to the genus level only, as well as dead trees. The y-axis represents the distribution density. Car, carotenoids; Chl *a*, Chlorophyll *a*; Chl *b*, Chlorophyll *b*; EWT, equivalent water thickness; LDMC, leaf dry matter content; LMA, leaf mass per area; SLA, specific leaf area.

## Discussion

### Model performance

In this study, we mapped multiple foliar functional traits in a mixed temperate forest using imaging spectroscopy combined with machine learning. We developed ensembles of PLSR models to estimate foliar traits from full-range reflectance spectra (400–2400 nm). Most of the models performed well with a validation R^2^ from 0.24 to 0.91 and %RMSE < 25 (except for carbon). Models achieved good to moderate performance for LMA, EWT, SLA, pigments (chl *a*, chl *b* and carotenoids), nitrogen and cellulose (Table 1). Models with poorest performance were observed for lignin, carbon, LDMC and hemicellulose. The validation R^2^ were lower for these traits. Still, most of the models show a good predictive accuracy as the %RMSE are within a decent value for this type of study (<25%, Table 1; Asner et al., 2015). The number of components used to build PLSR models ranged from 4 to 11, with nitrogen having the highest of all traits (11). However, this number is not considered high for fitting a canopy-level spectra-trait model (Wang et al., 2020).

Our first hypothesis, which stated that LMA, EWT and cellulose would be predicted with the highest accuracy, was partially validated. In fact, the best predictions were obtained for LMA, EWT and SLA. Carbon fractions (cellulose, hemicellulose, lignin) showed moderate to poor performance (Table 1). Estimating carbon fractions separately can be challenging (Homolová et al., 2013). In particular, canopy lignin is difficult to retrieve due to its similarity with other leaf structural compounds such as cellulose (Ustin et al., 2004), which has similar absorption features to lignin. Also, the low intrinsic trait variability of some traits can increase the challenge of estimating them accurately (Singh et al., 2015). For carbon and LDMC, the coefficient of variation was lower compared to other traits, indicating a small range in trait values (Table S1). LDMC showed poor performance, compared to leaf-level spectra-trait model (e.g., Kothari et al., 2023). The dry matter of leaves is a complex assemblage of organic compounds, and there are few studies that quantified it to the canopy-level using a PLSR approach. The relationship between canopy spectra and foliar traits are often lower in precision and accuracy than leaf-level models (Asner & Martin, 2008). In general, our models showed comparable results to those from other studies that used a similar PLSR approach, except for carbon that showed weaker results (Table S2). Overall, our study provides further evidence of the potential of imaging spectroscopy to accurately predict foliar traits, and also provides PLSR models that consider inter- and intraspecific variation and that can be applied to other temperate mixed forests in eastern Canada and elsewhere.

### Model interpretation

For model development, we used the full range VSWIR spectra (∼400–2400 nm). The VIP metric gives an insight into the relative importance of each wavelength to predict a foliar trait. When higher than 0.8, a VIP is considered significant (Singh et al., 2015). Figure 8 shows the VIP metric of the calibration models for each trait. Regions in the VIS, especially around 550 nm (green hump), were important to predict all traits. The red-edge (690–750 nm) showed significant VIPs for all traits, which is consistent with previous studies (Asner et al., 2015; Singh et al., 2015; Wang et al., 2020). Also, many wavelengths in the SWIR (e.g., 1527–1797 nm) had high VIPs for most traits.

**Figure 8.**
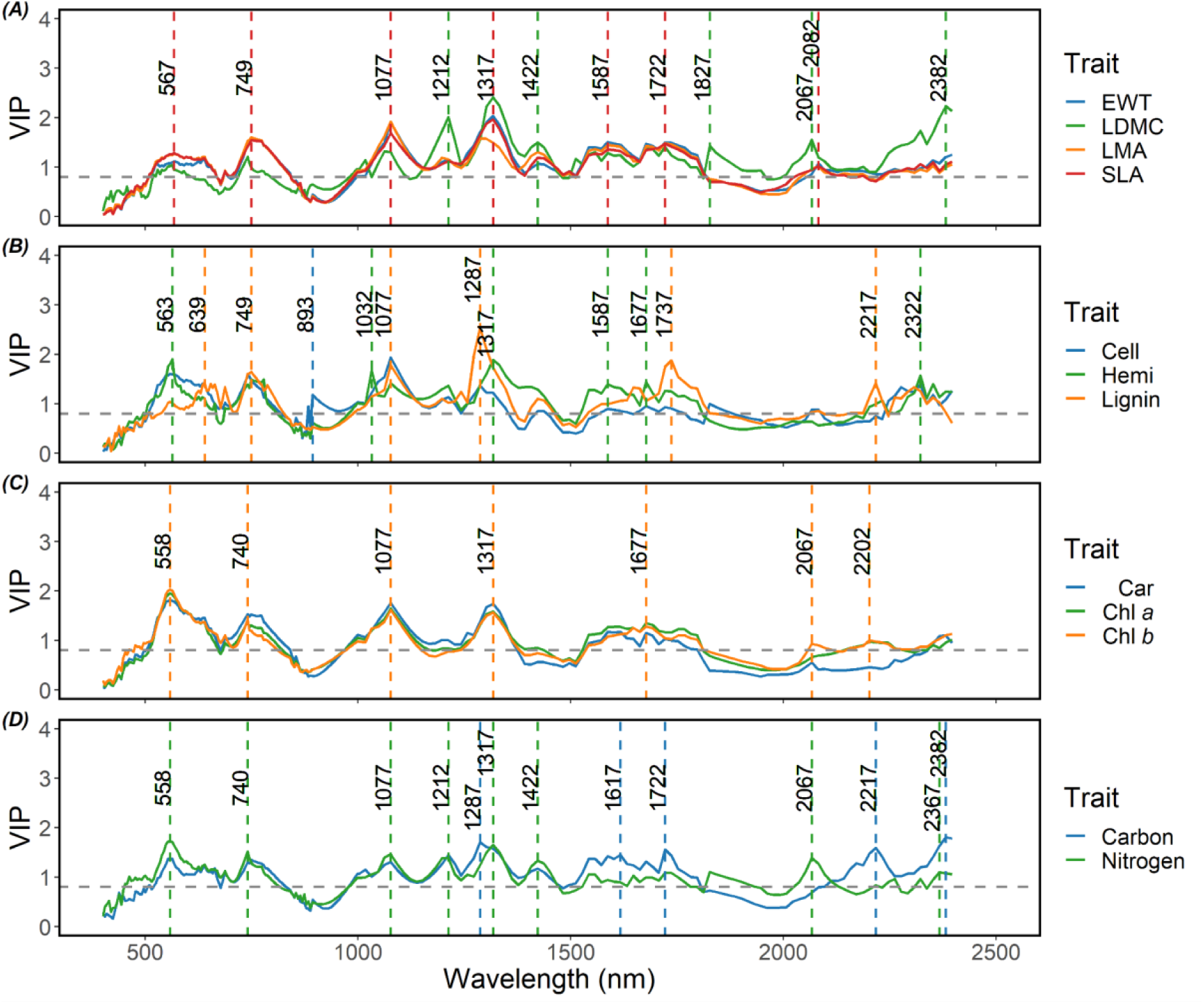
VIP metric from calibration models of all traits in the study. Traits shown are (A) equivalent water thickness (EWT), leaf dry matter content (LDMC), leaf mass per area (LMA), specific leaf area (SLA), (B) cellulose (Cell), hemicellulose (Hemi), lignin, (C) carotenoids (Car), chlorophyll *a* (Chl *a*), chlorophyll *b* (Chl *b*), (D) carbon and nitrogen. Horizontal gray dashed line represents a VIP of 0.8 and dashed lines are local maxima with their respective wavelength number in nm.

Important wavelengths for LMA and SLA were very similar throughout the spectrum, as they are inversely related to each other (Cornelissen et al., 2003). LDMC and EWT also showed a similar VIP pattern to that of LMA and SLA, but with higher peaks in the SWIR for LDMC. Important wavelengths for these traits corresponded to absorption features of water, cellulose, starch and lignin, chemicals that constitute dry mass of leaves (Curran, 1989; Wang et al., 2020). Carbon fractions (cellulose, hemicellulose and lignin) tended to show a similar pattern in VIP. Important wavelengths matched absorption features of cellulose, starch, sugar and lignin (Curran, 1989). All pigments (chlorophyll *a*, *b* and carotenoids) shared a similar VIP distribution, with important wavelengths matching known absorption features. Both chlorophylls showed VIP peaks that match with known absorption features in the VIS (558 nm) and the *red-edge* (740 nm; Blackburn, 2006). The VIP distribution for carbon and nitrogen differed mostly in the SWIR. Important wavelengths for nitrogen corresponded with known absorption features of proteins (e.g., 1020, 2060 and 2350 nm; Curran, 1989). Many absorption features interfere together due to trait covariance (Kokaly et al., 2009). For example, nitrogen showed local maxima related to N content in photosynthetic pigments (558, 740 nm). The similarities in VIP may also be related to trait covariance that influences the overall VIP metric, although it cannot alone explain it (Kothari et al., 2023). However, important wavelengths in models may not always be consistent with absorption features. Many of these features interfere together due to trait associations (Kokaly et al., 2009). Overall, important wavelengths in models were not always consistent with absorption features, and our second hypothesis was partially validated.

### Canopy trait distributions

Observing the distribution of leaf traits in certain areas can reveal some patterns that are consistent with our knowledge about foliar trait relationships. The leaf economic spectrum (LES) states that there is a trade-off in resource allocation in plants, resulting in species known as conservative and acquisitive (Wright et al., 2004). More conservative species tend to show a higher longevity, and generally show a higher leaf mass per area (LMA). On the contrary, more acquisitive species tend to grow faster and have a lower LMA. Figure 4 covers an area with a variety of species, both coniferous and broadleaf. Pixels of high LMA, low nitrogen and chlorophyll *a* correspond to conifers like *Abies balsamea*, *Pinus strobus*, or *Thuja occidentalis*. It is known that conifer species tend to exhibit traits on the conservative side of the LES (Díaz et al., 2016). This resource strategy means they generally have higher LMA and leaf thickness, as well as lower nutrients and photosynthetic rates than broadleaf species (Díaz et al., 2016; Zhang et al., 2020). This pattern was also observed in Figure S8 where individual trees are annotated at the species level.

The chemistry of aboveground organic inputs (e.g., fallen leaves) has a major influence on belowground processes like decomposition and nutrient cycling (Madritch et al., 2020). Figure 5 illustrates variation in foliar trait distribution across the transition to a poorly drained peatland. Species with higher LMA are generally associated with poor environments (Cornelissen et al., 2003). Water-saturated environments contain a high amount of organic matter, but few accessible nutrients for vegetation (Rodríguez-Iturbe & Porporato, 2007). This is consistent with the higher LMA that was observed across the peatland. High soil moisture creates anoxic conditions that reduce aerobic oxidation by bacteria, thus slowing decomposition rates. The availability of mineral nitrogen, the accessible form to plants, is limited in water-saturated environments due to slower decomposition (Rodríguez-Iturbe & Porporato, 2007). Nitrogen is linked to the amount of pigments since it is an essential compound in metabolic proteins such as RuBisCo (Evans, 1989). Since decomposition tends to be slower in water-saturated environments, this aligns with what was found in the peatland, i.e. low nitrogen and chlorophyll content. Cellulose, a complex compound hard to degrade, is also related to the rate of decomposition (Madritch et al., 2020). LDMC serves as an indicator of plants resources use, with cellulose representing a significant component of leaf dry matter (Garnier et al., 2001; Homolová et al., 2013). High LDMC species typically reflect a strategy to preserve nutrients (Garnier et al., 2001). This is consistent with the higher cellulose content found in the peatland, considering that fewer nutrients are available in high moisture soil (Rodríguez-Iturbe & Porporato, 2007).

Using annotated tree crowns from Cloutier et al. (2023), we compared the distribution of measured traits in the field and estimated traits corresponding to pixels within these polygons (Fig. 7). The predicted traits for these pixels mostly covered the range of traits measured during field sampling. The long right tail for certain traits may be attributed to a few higher values in the calibration dataset and the opposite can be suggested for carbon that showed a long left tail. The narrow peak in predicted traits suggest that the models capture less variability in species traits. This is confirmed with a lower standard deviation for all estimated traits (results not shown), compared to measured traits (Table S1). Canopy dominant species, such as *Acer rubrum* or *Betula papyrifera*, may be over-represented in the trait range. These two species were the most abundant in the tree polygon dataset (Cloutier et al., 2023). Overall, we feel that our dataset was large enough to capture the study site’s range of variation in canopy traits. While a few species were not included in the dataset (like *Fraxinus nigra* or *Prunus serotina*), these trees are relatively rare in the area and are unlikely to reach the canopy.

### Confidence of trait predictions

Mapping uncertainties related to trait predictions is essential for any potential ecological application of the trait maps. It can guide the user into what regions are predicted with low or high uncertainty. It also serves as insight for model performance in general (Wang et al., 2019). Our results showed high uncertainties in wetlands like swamps, fens and shallow waters, and vegetation near lake borders. Regions with high uncertainties can be explained by multiple factors: the difference in vegetation properties, mixed pixels, the presence of shallow water and canopy gaps. Indeed, grasses, forbs, shrubs, as well as wetland indicator plants, were not included in our dataset since we focused on trees. These types of vegetation, at the individual level, are smaller than the imagery’s pixel size (here 1.25 m). Consequently, this can result in mixed pixels, containing different plant lifeforms, and even soil and water.

The spectral signal of mixed pixels can be difficult to interpret in terms of trait predictions (Huete, 2004). For example, Hacker et al. (2022) showed that spectral mixing can affect predictive capacity of foliar traits and these effects vary depending on the trait of interest. As for shallow waters, vegetation covered by water creates differences in optical properties that are not captured by the models in the training task. Canopy gaps are more frequent near lake borders, which can explain the high uncertainties in these areas. It is known that compared to leaf-level reflectance, radiative properties of canopies are influenced by additional factors. It can be related either to the measuring instrument (ex. varying view and illumination angle) or to the target, like variation in leaf area index (LAI) and internal canopy shade (Feilhauer et al., 2010). Reducing the effect of canopy structure can help in the task of deriving leaf properties from reflectance spectra. In this study, we applied an NDVI filter and a shadow threshold to remove most of the non-vegetated and shadowed pixels. An NDVI filter is useful to remove low LAI pixels, but gaps and shadows may need to be handled with more than a shadow threshold in the context of trait mapping over large areas (Asner et al., 2015). LiDAR has been proved useful in previous studies to pre-screen imagery, identifying gaps and shadows (Asner & Martin, 2008; Chadwick & Asner, 2016; Martin et al., 2018), but this approach was not applied in our study.

Relative uncertainties expressed as a percentage allow us to compare uncertainties in predictions of foliar traits with different units. Over 90% of pixels that fell in the predefined range from field measurements (see section *Foliar trait maps*) had relative uncertainties lower than 25% for all traits except for LMA (Fig. S7). This trait showed the lowest percent of high-confidence pixels (87%), which may seem unusual given its higher accuracy across all models (Table 1). One could expect that LMA would have the highest percent of high-confidence pixels since the model has the lowest uncertainties. This may be linked to the higher negative values compared to other traits (∼0.81%) that were removed from analysis, decreasing the number of total pixels for the percentage calculation. Overall, pixels with high confidence values were always higher than 85%. These results support the confidence in the trait maps’ reliability.

### BRDF effects

Some trait maps showed visual artifacts that appear as a long stripe between pixels on the right and left sides of a flightline (Fig. S5). For example, LDMC, nitrogen and hemicellulose models showed more artifacts than other traits in the first flightline. This suggests that the presence of artifacts is not necessarily related to the model’s overall prediction performance (Table 1). In this case, it appears to be the result of bidirectional reflectance distribution function (BRDF) effects. The BRDF describes how light is reflected by a surface depending on the solar and view geometries. Unwanted gradients of brightness can arise from BRDF effects between flightlines of imagery. This is generally rectified with a BRDF correction (Queally et al., 2022). However, such corrections can be challenging in some contexts. In fact, one of the imaging spectrometers used in this study, SASI-600, is composed of two sensors whose data are merged as a single image during preprocessing. Consequently, the SASI dataset showed a subtle difference in brightness between the left and right side pixels in a flightline. Even with the standard normalization applied here (see section *Imaging spectroscopy data*), the effects were reduced for some wavelengths better than others but were not completely eliminated. These artifacts may be more apparent for models that were strongly influenced by wavelengths with greater BRDF effects.

To investigate this further, we ran a principal component analysis (PCA) on a subset of pixels from the first flightline, containing pixels from both the right and left sides of the SASI dataset. Among the first nine principal components (PCs), PC2 and PC4 showed the strongest visual artifacts (Fig. S9). Absolute values of loadings for PC2 and PC4 showed that particular wavelengths in the near infrared (NIR, ∼800–1300 nm) had high loadings. This was also the case for wavelengths at the end of SWIR2 (∼2000–2500 nm). Thus, they contributed highly to the formation of PC2 and PC4. Also, these bands seemed to correspond with high VIP of trait models (Fig. S10). Specifically, some of the loadings’ local maxima were close to wavelengths important for estimating LDMC (1212, 1422, 1828, 2398 nm) nitrogen (1318, 1422, 2068 nm) and hemicellulose (1318, 1512, 1677 nm). If the model heavily relied on these bands affected by BRDF effects, this may explain the pronounced visual artifacts in the LDMC, nitrogen and hemicellulose maps of the first flightline. This could have been corrected with a radiometric alignment, but this was beyond the scope of this study. This finding emphasizes the importance for precise and accurate corrections of BRDF effects in large-scale studies of mapping foliar functional traits.

## Conclusion

In summary, we generated maps of 12 different foliar traits in a mixed temperate forest and their uncertainties, based on a method combining imaging spectroscopy and PLSR modeling. A unique feature of our study is that the foliar trait measurements used to train the models came from individual trees which could be accurately georeferenced to the imaging spectroscopy data, thus ensuring that intraspecific trait and spectral variability could be represented in our models. Our models’ performances varied among traits, with LMA, EWT and SLA showing the best performance. Almost all trait models had uncertainties that are considered adequate with this type of study. All trait maps will soon be available in the Federated Research Data Repository (FRDR), along with all model coefficients. The accessibility of our data and models promotes a collaborative approach to advance our understanding of foliar traits, on which plant functional ecology relies. Studies like ours that work at a high spatial resolution can serve as validation data for hyperspectral satellite missions (Wang et al., 2020). Furthermore, they can provide a baseline for further ecological research in the context of functional biodiversity, such as inter-specific variation of foliar traits. With CNN (convolutional neural network) models now able to identify pixels to the species levels (e.g., Cloutier et al., 2023), it opens the possibility to study inter-specific variation of foliar traits. This can greatly enhance our understanding of ecosystems functional properties. This work adds to the extensive research aiming to use remote sensing to assess and monitor forest biodiversity at larger scales (Cavender-Bares et al., 2020).

## Acknowledgements

This study was conducted as part of the Canadian Airborne Biodiversity Observatory (CABO) which was funded by a Discovery Frontiers grant (NSERC; #5091902017) from the Natural Sciences and Engineering Research Council of Canada (NSERC). We thank Juan Pablo Arroyo-Mora (National Research Council of Canada, Flight Research Lab; NRC-FRL) who acquired airborne imagery, and Deep Inamdar (Applied Remote Sensing Lab, McGill University; ARSL) for the preprocessing of airborne imagery. We thank Myriam Cloutier, Sabrina Demers-Thibeault, Étienne Morissette, Charles Picard-Krashevski and Ariane Roberge for their help during the field campaign. We thank the Station de Biologie des Laurentides (SBL) of Université de Montréal for facilitating the field sampling. We thank Shan Kothari for his feedback on model development and Christine Wallis for her insights on image post-processing.

## Author contributions

AG and EL designed the study and acquired the trait data. AG analyzed and interpreted the data, and wrote the first draft of the manuscript. All authors participated in the revision of the paper.

## Data availability

The tree dataset (field-measured foliar traits, mean reflectance spectra), model coefficients and trait maps are available in the <u>LEFO collection</u> of the Federated Research Data Repository (FRDR, https://doi.org/10.20383/103.0922). All scripts, except for the pre-processing of the imaging spectroscopy, can be found as a GitHub repository (https://github.com/alicegravel98/Trait-mapping-SBL.git).

