## Supplementary Materials for "Mapping canopy foliar functional traits in a mixed temperate forest using imaging spectroscopy"

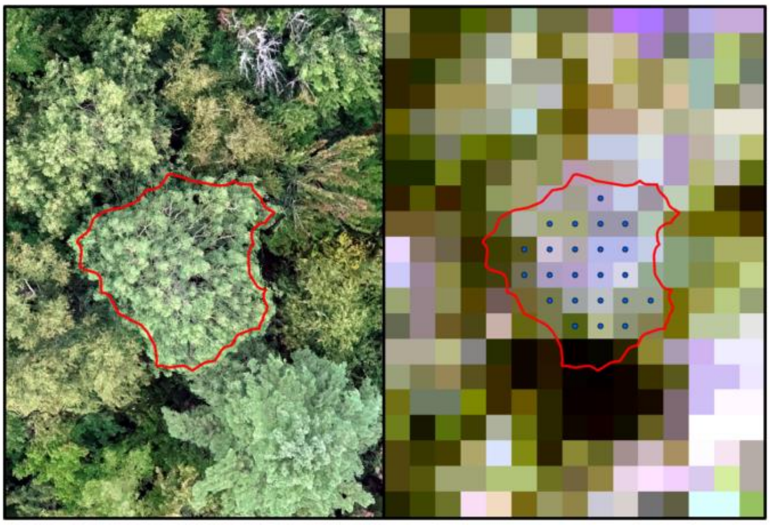

**Figure S1. Example of pixels extracted to average spectra within a tree crown.** On the left, a tree crown is presented with a 0.5 inner buffer in red. On the right, hyperspectral pixels extracted within the buffer to obtain one mean reflectance are presented as blue dots.

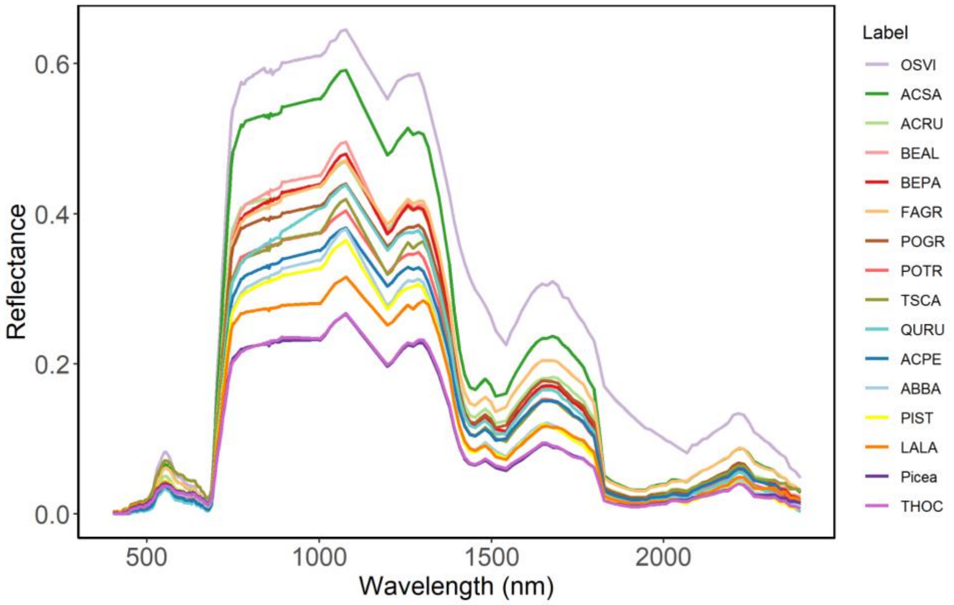


**Figure S2. Smoothed mean canopy reflectance by species.** Species labels are listed in descending order of reflectance. OSVI, *Ostrya virginiana*; ACSA, *Acer saccharum*; ACRU, *Acer rubrum*; BEPA, *Betula alleghaniensis*; FAGR, *Fagus grandifolia*; POGR, *Populus grandidentata*; POTR, *Populus tremuloides*; TSCA, *Tsuga canadensis*; QURU, *Quercus rubra*; ACPE, *Acer pensylvanicum*; ABBA, *Abies balsamea*; PIST, *Pinus strobus*; LALA, *Larix laricina*; Picea, *Picea sp.*; THOC, *Thuya occidentalis*.

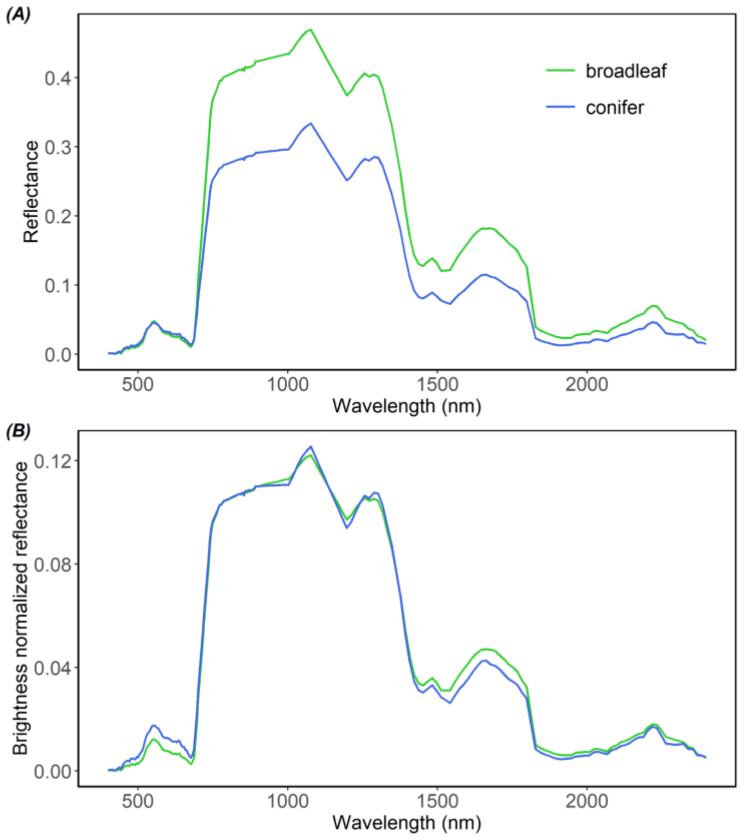


**Figure S3. Mean canopy reflectance (A) and mean brightness normalized reflectance (B) of tree crowns extracted from the imagery by vegetation type.** Broadleaf species are in green and conifer species are in blue.

**Table S1. Summary statistics of foliar traits from sampled tree crowns used in calibration and validation (n = 166).** SD, standard deviation; CV, coefficient of variation. Chl *a*, chlorophyll *a*; Chl *b*, chlorophyll *b*; EWT, equivalent water thickness; LDMC, leaf dry matter content; LMA, leaf mass per area; SLA, specific leaf area.

| Traits | units | min | max | mean | SD | CV |
|---|---|---|---|---|---|---|
| Carbon | % | 45.58 | 51.19 | 48.77 | 1.03 | 2.11 |
| Carotenoids | mg $g^{-1}$ | 0.10 | 0.83 | 0.38 | 0.15 | 39.47 |
| Cellulose | % | 6.19 | 18.73 | 10.21 | 2.77 | 27.13 |
| Chl *a* | mg $g^{-1}$ | 0.38 | 3.58 | 1.58 | 0.66 | 41.77 |
| Chl *b* | mg $g^{-1}$ | 0.12 | 1.00 | 0.48 | 0.20 | 41.67 |
| EWT | mm | 0.06 | 0.35 | 0.15 | 0.07 | 46.67 |
| Hemicellulose | % | 6.15 | 28.33 | 13.70 | 4.45 | 32.48 |
| LDMC | mg $g^{-1}$ | 327.20 | 546.68 | 455.52 | 44.23 | 9.71 |
| Lignin | % | 4.51 | 19.37 | 10.44 | 2.94 | 28.16 |
| LMA | g $m^{-2}$ | 49.87 | 433.44 | 142.91 | 80.73 | 56.49 |
| Nitrogen | % | 0.69 | 3.06 | 1.60 | 0.47 | 29.37 |
| SLA | $m^2$ $kg^{-1}$ | 2.31 | 20.05 | 8.91 | 3.74 | 41.98 |

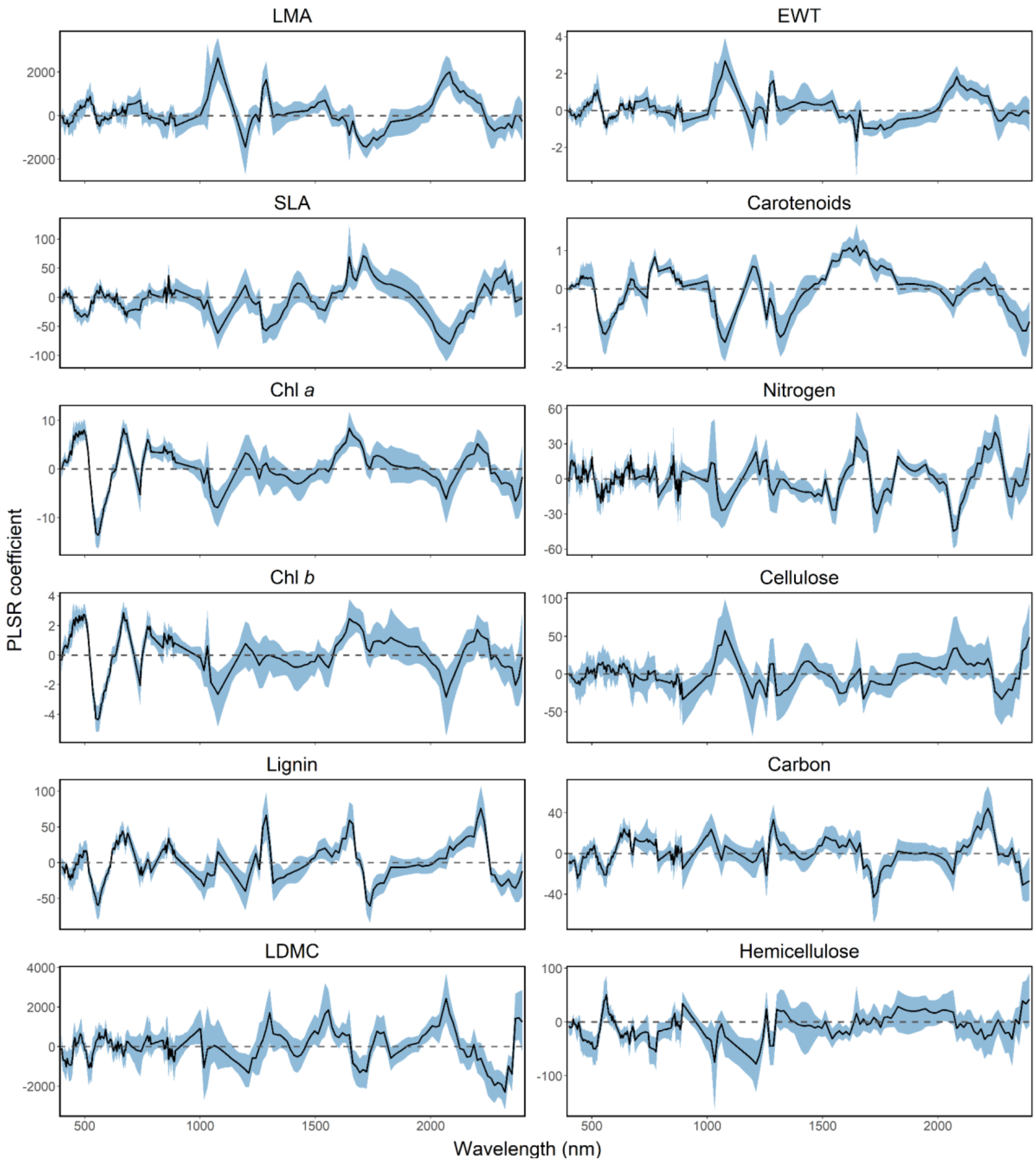


**Figure S4. Jackknife regression coefficients for 100 iterations.** The black line shows the mean spectral weighting vector, and the blue area represents the 95% confidence interval. The dashed gray line indicates a coefficient of zero. Coefficient close to zero reflects a low contribution of that wavelength to the prediction of the trait of interest. LMA, leaf mass per area; EWT, equivalent water thickness; SLA, specific leaf area; Chl *a*, chlorophyll *a*; Chl *b*, Chlorophyll *b*; LDMC, leaf dry matter content.

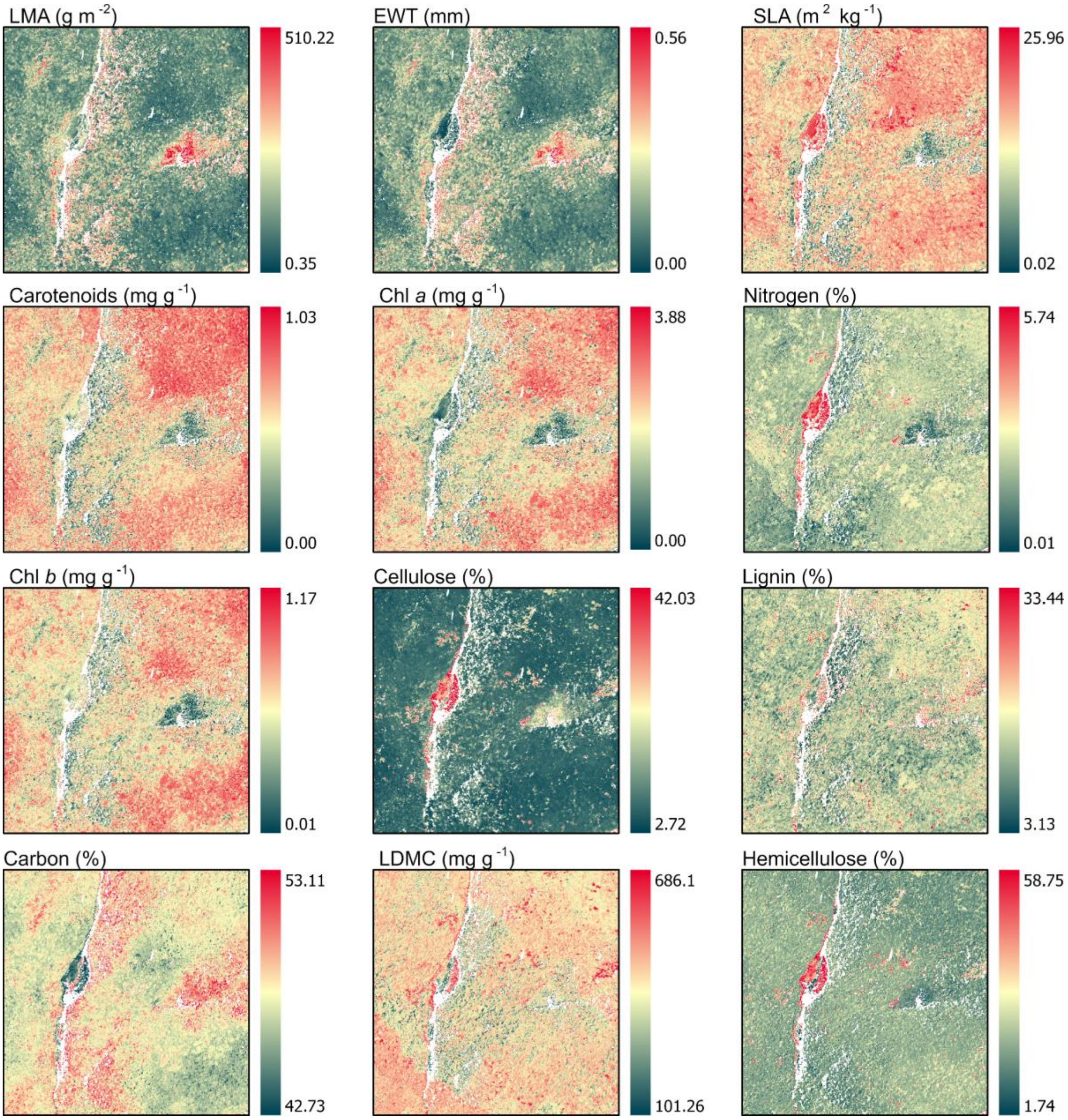


**Figure S5. Panels of predicted foliar traits, obtained by applying an ensemble of PLSR models to the imaging spectroscopy data.** LMA, leaf mass per area; EWT, equivalent water thickness; SLA, specific leaf area; Chl *a*, chlorophyll *a*; Chl *b*, chlorophyll *b*; LDMC, leaf dry matter content.

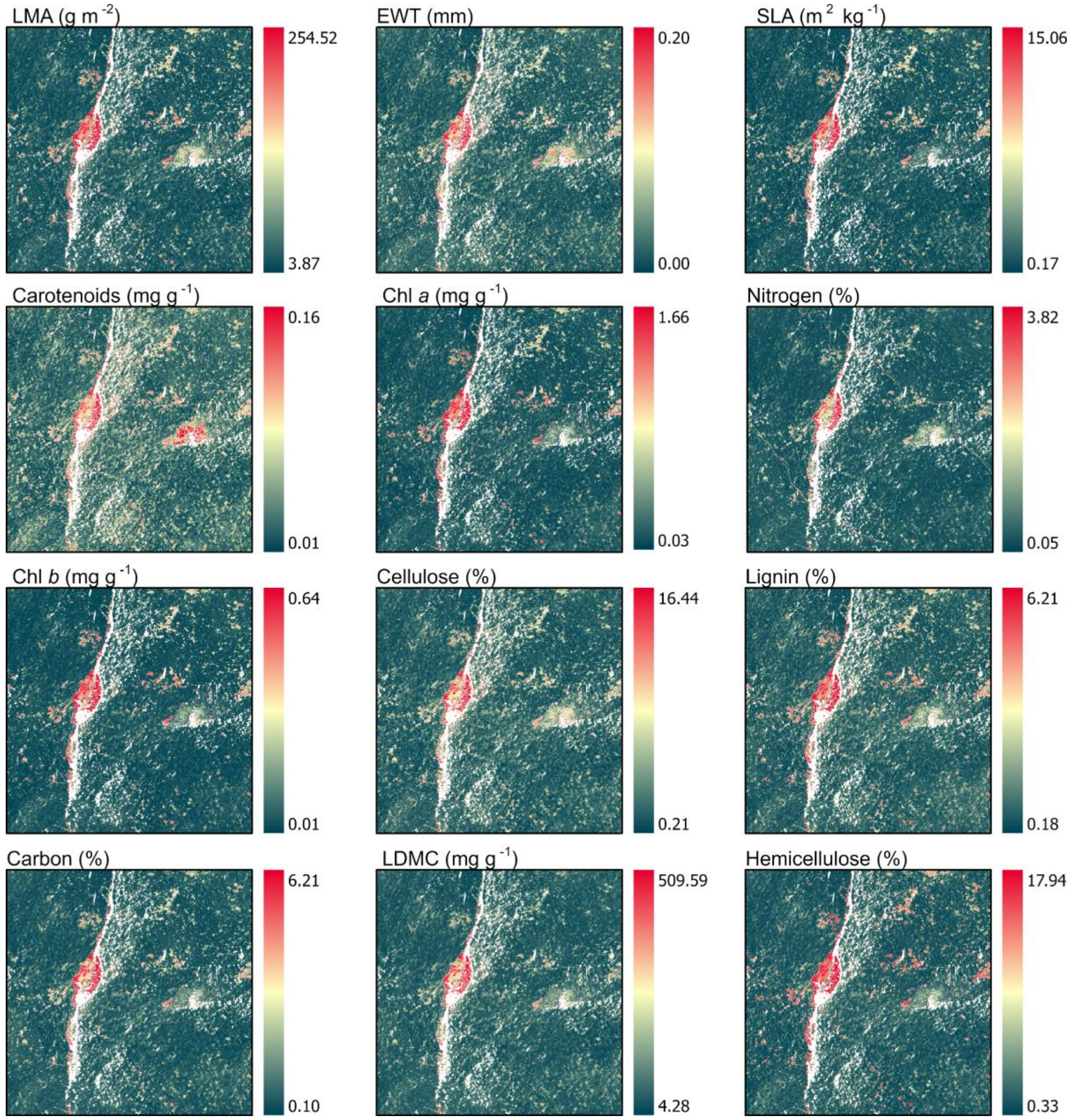


**Figure S6. Panels of foliar traits uncertainties, obtained by calculating SD of 100 permutations.** High uncertainties are found near lake borders and shallow water. LMA, leaf mass per area; EWT, equivalent water thickness; SLA, specific leaf area; Chl *a*, chlorophyll *a*; Chl *b*, chlorophyll *b*; LDMC, leaf dry matter content.

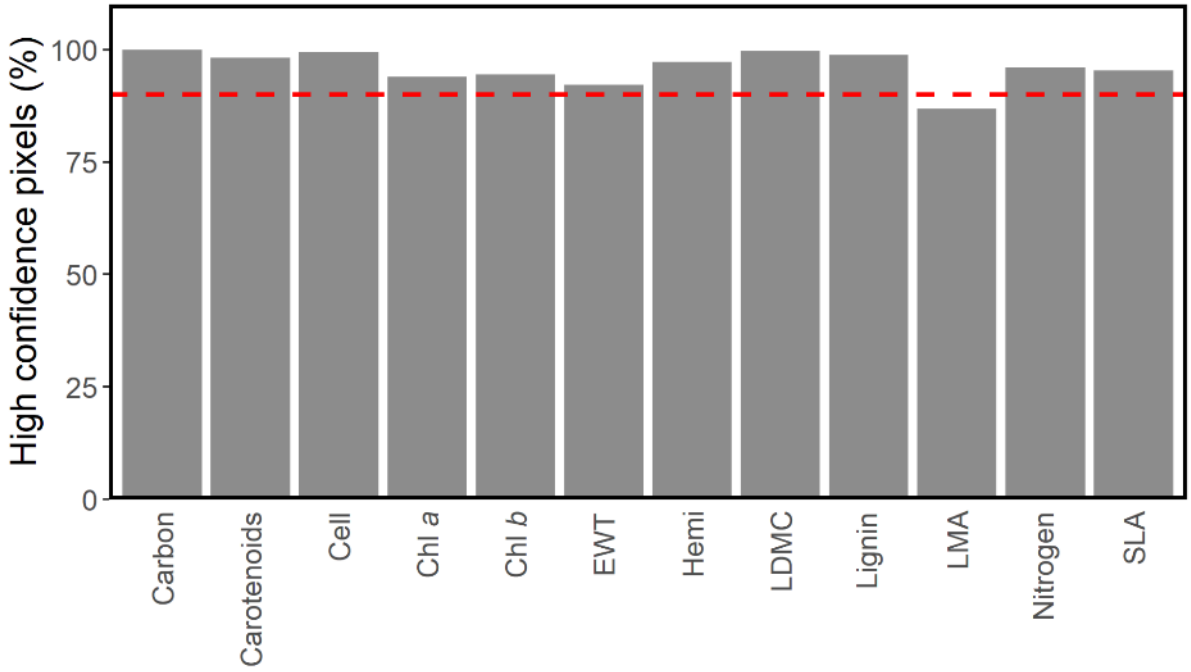


**Figure S7. Percent of pixels with high confidence for each trait and flightlines combined.** Dashed red line indicates 90% pixels with high confidence. Cell, cellulose; Chl *a*, chlorophyll *a*; Chl *b*, chlorophyll *b*; EWT, equivalent water thickness; Hemi, hemicellulose; LDMC, leaf dry matter content; LMA, leaf mass per area; SLA, specific leaf area.

**Table S2. Comparison of model predictive accuracies between this study and other studies using PLSR approach (Asner et al., 2015; Martin et al., 2018; Wang et al., 2020).** Gray cells indicate no model was developed for the trait. The asterisk indicates that the statistics of other studies for this trait are for models combining both chlorophylls (*a* and *b*). Chl *a*, chlorophyll *a*; Chl *b*, chlorophyll *b*; EWT, equivalent water thickness; LDMC, leaf dry matter content; LMA, leaf mass per area; SLA, specific leaf area.

| Traits | **This study** (temperate forest) | | | **Asner et al. 2015** (tropical forest) | | | **Martin et al. 2018** (tropical forest) | | | **Wang et al. 2020** (temperate, subtropical forest and grasslands) | | |
|---|---|---|---|---|---|---|---|---|---|---|---|---|
| | **Units** | **$R^2$** | **%RMSE** | **Units** | **$R^2$** | **%RMSE** | **Units** | **$R^2$** | **%RMSE** | **Units** | **$R^2$** | **%RMSE** |
| Carbon | % | 0.34 | 26.13 | % | 0.44 | 6.09 | % | 0.41 | 4.70 | mg $g^{-1}$ | 0.69 | 12.44 |
| Carotenoids | mg $g^{-1}$ | 0.68 | 17.83 | mg $g^{-1}$ | 0.49 | 21.20 | | | | mg $g^{-1}$ | 0.52 | 14.58 |
| Cellulose | % | 0.53 | 21.31 | % | 0.20 | 21.04 | % | 0.18 | 32.30 | mg $g^{-1}$ | 0.61 | 11.69 |
| Chl *a** | mg $g^{-1}$ | 0.66 | 17.47 | mg $g^{-1}$ | 0.58 | 28.14 | mg $g^{-1}$ | 0.21 | 29.80 | mg $g^{-1}$ | 0.60 | 13.22 |
| Chl *b* | mg $g^{-1}$ | 0.64 | 17.24 | | | | | | | | | |
| EWT | mm | 0.87 | 11.88 | | | | | | | g $m^{-2}$ | 0.73 | 9.70 |
| Hemicellulose | % | 0.24 | 23.33 | | | | | | | | | |
| LDMC | mg $g^{-1}$ | 0.32 | 23.79 | | | | | | | | | |
| Lignin | % | 0.44 | 20.67 | % | 0.26 | 24.08 | % | 0.22 | 32.40 | mg $g^{-1}$ | 0.47 | 17.97 |
| LMA | g $m^{-2}$ | 0.91 | 9.59 | g $m^{-2}$ | 0.53 | 19.25 | g $m^{-2}$ | 0.81 | 21.60 | g $m^{-2}$ | 0.77 | 9.24 |
| Nitrogen | % | 0.65 | 18.42 | % | 0.48 | 15.19 | % | 0.54 | 24.40 | mg $g^{-1}$ | 0.61 | 14.38 |
| SLA | $m^2$ $kg^{-1}$ | 0.82 | 13.35 | | | | | | | mm $mg^{-1}$ | 0.66 | 11.94 |

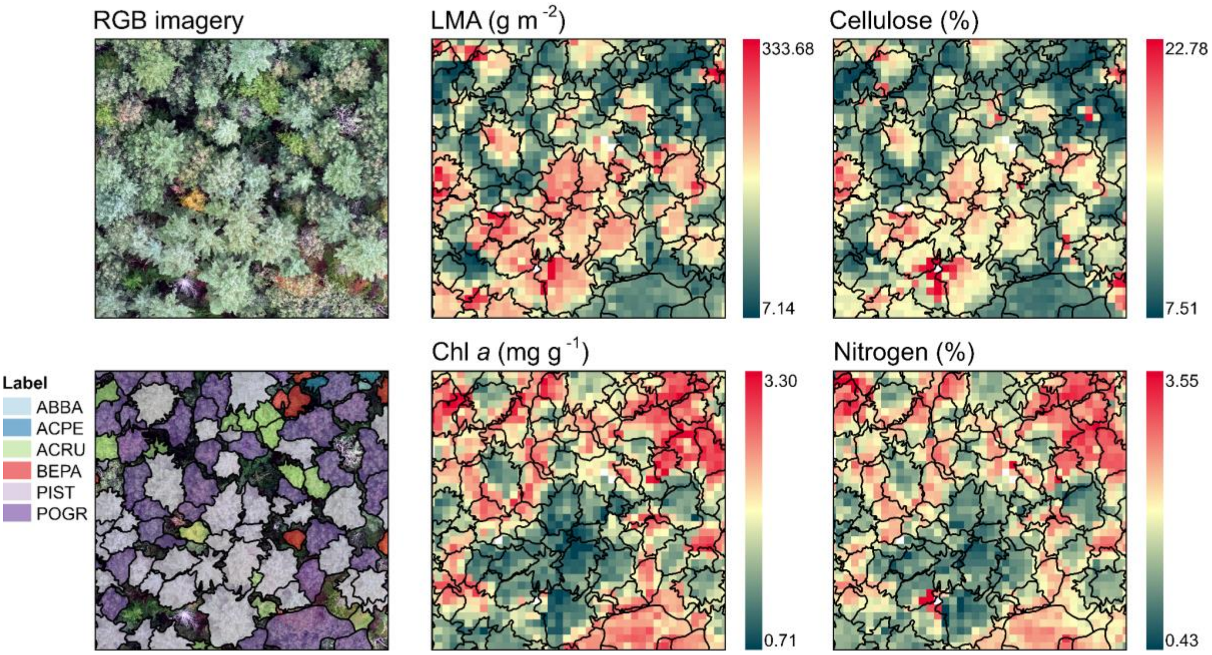


**Figure S8. Maps of four different foliar traits and RGB imagery and individual trees annotated at the species level (LMA, leaf mass per area; cellulose; Chl *a*, chlorophyll *a* and nitrogen).** Species are represented in different colors in RGB imagery (Cloutier et al., 2023). Black outlined polygons in trait maps represent the same trees.

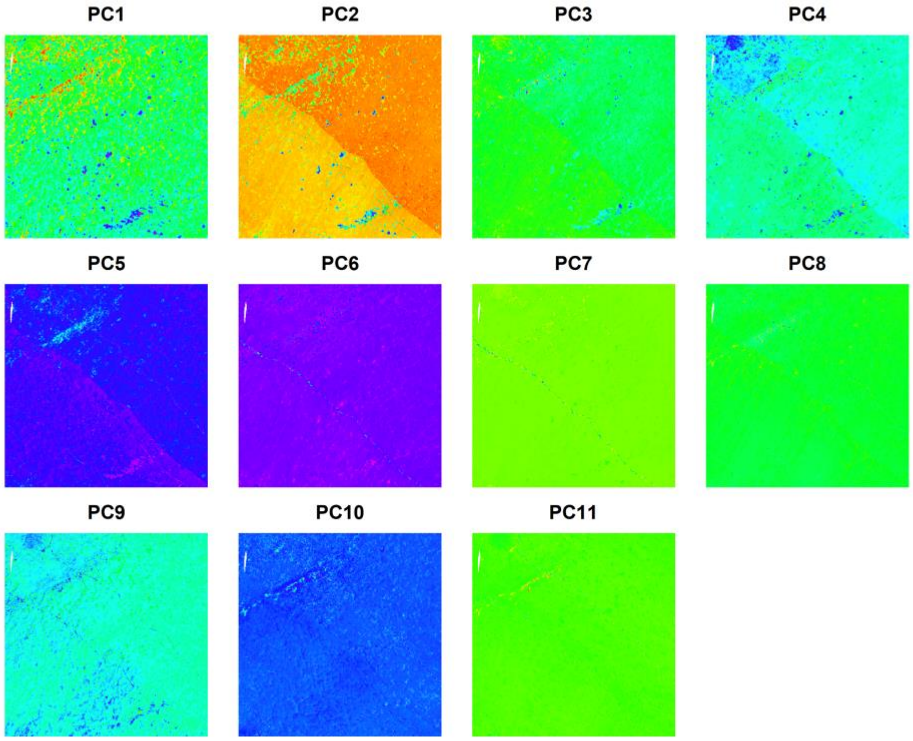


**Figure S9. The first 9 principal components (PCs) derived from the SASI dataset image cube.** Together, they represent 99.1% of the spectral variation in selected pixels from the first flightline. Artifacts related to BDRF effects are mostly visible in PC2 and PC4, where pixels on the right and left sides of the imagery have brightness differences.

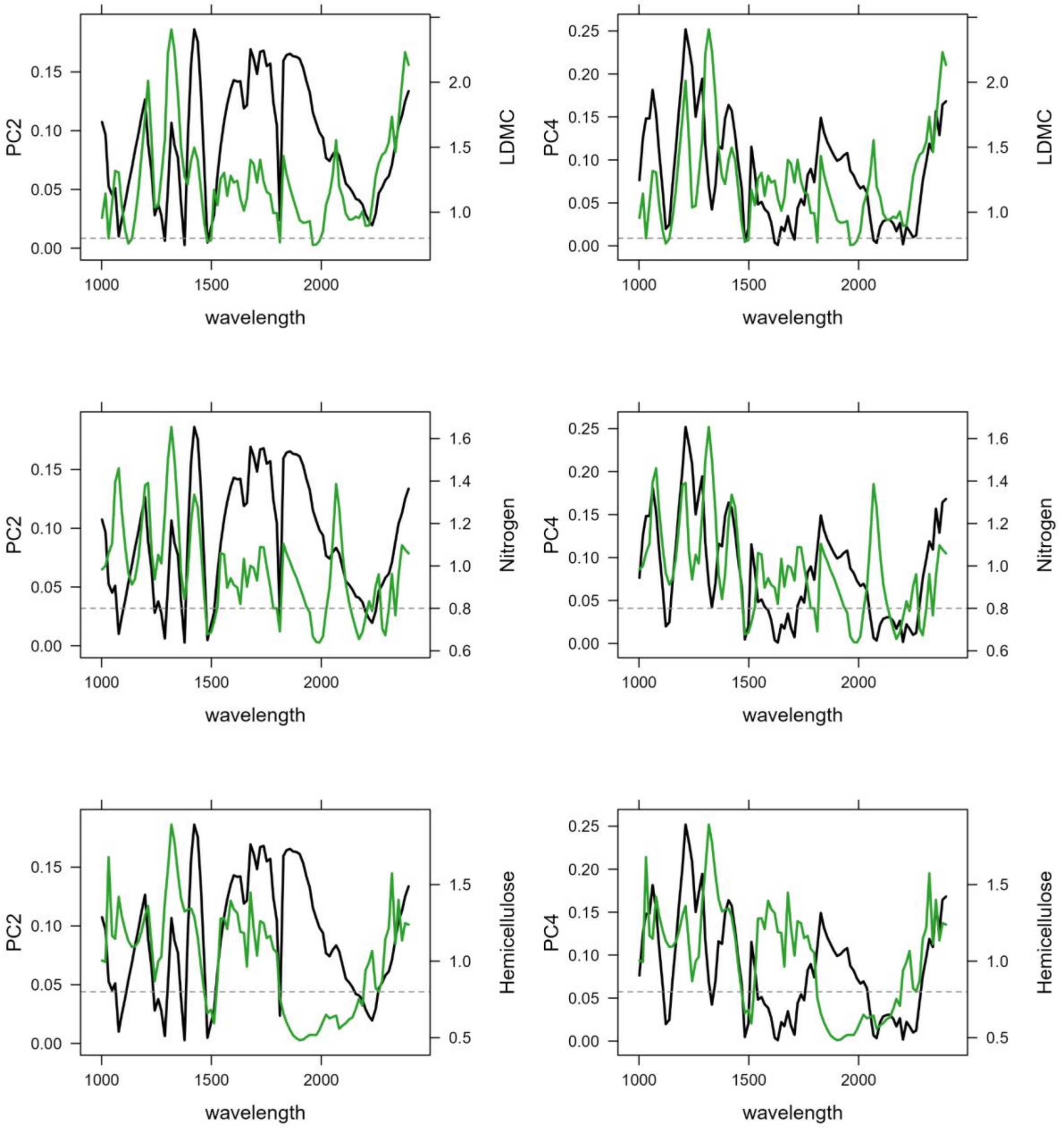


**Figure S10. Principal components (PCs) 2 and 4 against variable importance in projection (VIP) for traits mostly affected by BRDF effects (LDMC, leaf dry matter content; nitrogen and hemicellulose).** VIPs are shown in green and loadings in black. Horizontal gray dashed line represents a VIP of 0.8.
